# Imaging the spontaneous flow of thought: Distinct periods of cognition contribute to dynamic functional connectivity during rest

**DOI:** 10.1101/527804

**Authors:** Javier Gonzalez-Castillo, César Caballero-Gaudes, Natasha Topolski, Daniel A. Handwerker, Francisco Pereira, Peter A. Bandettini

## Abstract

Brain functional connectivity (FC) changes have been measured across seconds using fMRI. This is true for both rest and task scenarios. Moreover, it is well accepted that task engagement alters FC, and that dynamic estimates of FC during and before task events can help predict their nature and performance. Yet, when it comes to dynamic FC (dFC) during rest, there is no consensus about its origin or significance. Some argue that rest dFC reflects fluctuations in on-going cognition, or is a manifestation of intrinsic brain maintenance mechanisms, which could have predictive clinical value. Conversely, others have concluded that rest dFC is mostly the result of sampling variability, head motion or fluctuating sleep states. Here, we present novel analyses suggesting that rest dFC is influenced by short periods of distinct mental processing, and that the cognitive nature of such mental processes can be inferred blindly from the data. As such, several different behaviorally relevant whole-brain FC configurations may occur during a single rest scan even when subjects were continuously awake and displayed minimal motion. In addition, using low dimensional embeddings as visualization aids, we show how FC states—commonly used to summarize and interpret resting dFC—can accurately and robustly reveal periods of externally imposed tasks; however, they may be less effective in capturing periods of distinct cognition during rest.

## INTRODUCTION

As the field of functional Magnetic Resonance Imaging (fMRI) moves away from group-level reports and becomes increasingly interested in the types of single-subject evaluations that are necessary for clinical diagnosis and prognosis; there is an increasing focus on examining how brain states change over time rather than treating each scan as a static snapshot of a person’s brain function. For example, individual scans today are often described in terms of a limited set of recurring, short-duration (tens of seconds), whole-brain FC configurations named FC states (1–3). Metrics describing their dwell times, ordering and frequency of transitions can then be used to quantify different aspects of empirically observed dFC (4). Many questions remain both about the etiology of empirically observed systems-level FC dynamics; as well as regarding the ability of models to accurately capture behavioral, cognitive and clinically relevant dynamic phenomena.

One well accepted principle is that task performance can modulate dFC across the brain (5), even if the brain never abandons an overall small-world configuration (6). As humans perceive external stimuli and engage with cognitive tasks, patterns of communication across brain networks reshape (7), modularity decreases (8), communication hubs relocate (9), and overall variability of FC estimates drops (10). FC modulations induced by tasks are sufficiently profound as to allow prediction of subsequent perceptual outcomes such as response time (11) and pain levels (12). Furthermore, FC estimates computed from only seconds long portions of data can be used to classify scans according to tasks (13) and to segment multi-task scans into task-homogenous periods (3).

Nonetheless, despite empirical reports of dFC in resting humans (1), macaques (14) and rodents (15), a similar level of consensus does not exist regarding the significance of dFC phenomena while at rest—with rest referring to a subject’s state when instructed to stay awake and not required to perform any given task or pay attention to any specific external stimuli. Those who hypothesize rest dFC to be neuronally relevant have explored the phenomenon in the context of consciousness (16), development (17) and clinical disorders (2, 18, 19). These studies have shown how the complexity of dFC decreases as consciousness levels decline (16), how dynamic inter-regional interactions can be used to predict brain maturity (17), and how dFC derivatives (e.g., dwell times) can be diagnostically informative for conditions such as schizophrenia (2), mild cognitive impairment (18), and autism (19). Yet, many others have raised valid concerns regarding the ability of current dFC estimation methods to capture neuronally relevant dFC at rest (20–23). These concerns include a lack of appropriate null models to discern real dynamics from sampling variability (20), improper pre-processing leading to spurious dynamics (21), and excessive temporal smoothing that hinder our ability to capture sharp and rapid transitions of interest (a real concern for sliding window techniques used to estimate FC states; (22, 24)). Finally, some have stated that resting dFC is primarily a manifestation of sampling variability, residual head motion artifacts, and fluctuations in sleep state; and as such, mostly irrelevant (25).

One cause of such discrepant views is that it is challenging to demonstrate the potential cognitive correlates of resting dFC; especially given the unconstrained cognitive nature of rest and scarcity of methods to reliably infer the cognitive correlates of whole-brain FC patterns. When subjects are instructed to quietly rest, retrospective reports demonstrate that subjects often engage in a succession of self-paced cognitive processes including inner speech, musical experience, visual imagery, episodic memory recall, future planning, mental manipulation of numbers, and periods of heightened somatosensory sensations (26). Reconfigurations of FC patterns during rest could, to some extent, be a manifestation of this flow of covert self-paced conscious cognition (16); even if other factors (e.g., random exploration of cognitive architectures (27), fluctuations in autonomic system activity (28), self-initiated body motion (29) and arousal levels (25)) also contribute. Here we present a series of experiments aimed at exploring this hypothesis, namely that distinct periods of covert cognition are significant contributors to observable resting dFC. In parallel, we also evaluate the ability of FC states— one of the most prominent approaches to summarize rest dFC—to capture those hypothesized periods of distinct cognition during rest.

To explain these diverse and sometimes contradictory observations regarding dFC, we extend current FC state methodology in two ways. First, we combine hemodynamic deconvolution (30) and activity-based reverse-inference (31) to map FC states onto cognitive states. An FC state is commonly defined in terms of a representative FC matrix and a timeline (when it occurs). The deconvolution step allows us to also generate a representative “activity” map per FC state. Those FC state “activity” maps are subsequently input to a pre-existing decoding framework (*Neurosynth*; (31)) able to map whole-brain activity patterns into ranked lists of cognitive processes likely associated with the input map. A direct mapping from representative FC matrices to cognitive processes might be preferable but, unfortunately as of today, such systems do not exist. In the past, hemodynamic deconvolution and reverse-inference have proven successful at decoding the nature of self-driven body motion during rest scans (29). Here, we extend that approach for the purpose of attaining open-ended cognitive decoding of FC states. The second way in which we extend current FC state methodology is the use of manifold learning techniques to generate low dimensional representations of time-varying FC that help visualize how connectivity evolves and help identify periods of distinct cognition during rest.

The proposed methods are first evaluated on a multi-task^1^ fMRI dataset (3) for which precise information about the timing and nature of cognitive states (as dictated by task demands) is available. Next, we apply the same methods to 15 mins long resting state scans with low head motion and infrequent eye closures from the human connectome project (HCP, (32)). This way, we examine the possible presence of periods of distinct cognition during rest, and the ability of FC states to uncover such periods. Comparative analyses across both scenarios (multi-task and rest) are important as substantial differences in the timing and nature of externally driven versus self-paced cognitive processes, and their manifestations in terms of dFC phenomena, may modulate the ability of models to capture cognitively relevant information across both scenarios.

Our results confirm both that FC states can accurately capture periods of distinct cognition driven by external task demands, as previously shown (3), and also that the proposed extension for the FC state framework permits accurate inference of the cognitive processes underlying each task FC state. Next, we show that although periods of distinct cognition are present during pure rest, and their cognitive correlates can be inferred in a similar manner, FC state modeling—at least in the specific form examined here—does not identify discrete cognitive states as consistently as with the task data. Overall, our work suggests that resting dFC is influenced by ongoing cognition, and therefore more than a single correlation structure may be necessary to entirely describe rest scans. At the same time, our results suggest that FC states do not consistently capture periods of distinct cognition during rest, and that their estimation and interpretation as a model for rest dFC should be updated accordingly.

## RESULTS

### FC States and Cognition in Multi-task Scenario

After pre-processing, FC states were estimated following procedures previously described in (3) for the multi-task dataset, which include: timeseries extraction for the 200-ROI Craddock Atlas (33), dimensionality reduction via PCA, estimation of sliding window correlation (window duration/step= 30/1.5s), Fisher’s Z transformation, and connectivity matrix vectorization (Suppl. Fig. 1). This procedure resulted in 998 connectivity vectors (or “snapshots”) per run, where each vector element is an estimation of pair-wise windowed connectivity. Next, snapshots were clustered into four groups (the number of different tasks) using *k*-*means* (34). Each resulting group is an FC state described in terms of a representative FC matrix and a timeline of windows contributing to the state. Agreement between task and FC states timing was quantified via the Adjusted Rand Index (ARI; (35))^2^. We performed these analyses under two different scenarios: (a) using only task homogenous windows—namely those that fall completely within the temporal span of individual task blocks (as was previously published (3)); and (b) using all available windows, which also include windows spanning more than one task block. The goal of comparing these two scenarios is to evaluate to what extent having cognitively inhomogeneous windows may affect our ability to correctly segment the multi-task scans. On average (see Suppl. Fig. 2), FC states faithfully recovered task timing in both scenarios: *ARI* = 0.89 ± 0.18 for the case when all windows entered the analyses; and *ARI* = 0.97 ± 0.10 when only task homogenous windows are considered. Yet, there was a significant difference in ARI across scenarios (*T*=2.25; *p*=0.02). For the “*all windows*” scenario, which is the primary focus here, 5 subjects out of 20 had an ARI < 0.8 (moderate or poor).

**Figure 1.**
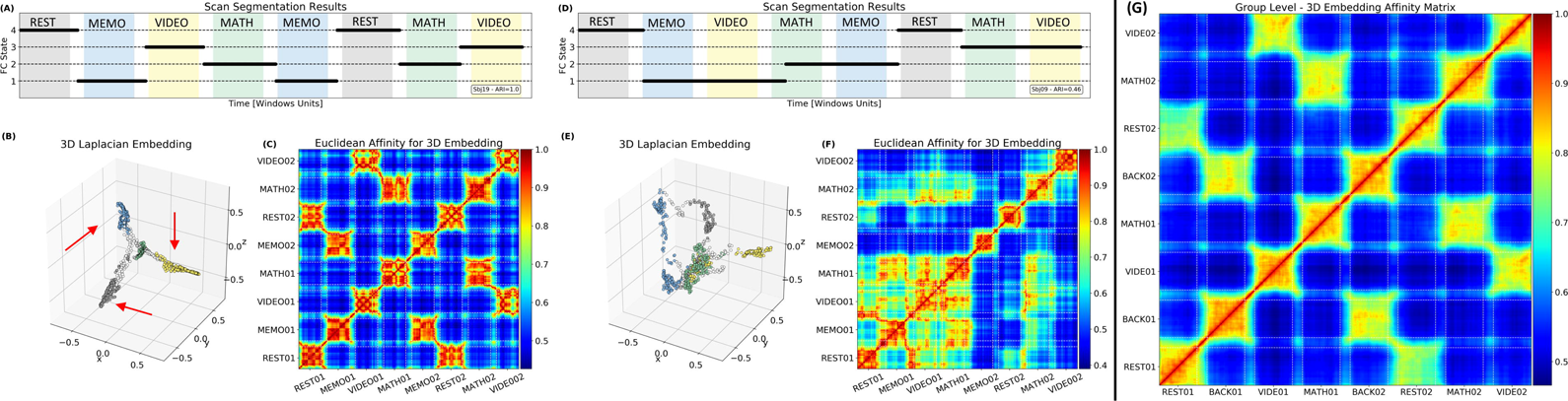
(A) FC state timeline, (B) 3D embedding, and (C) associated affinity matrix for one multi-task subject that showed good agreement (ARI = 1.0) between FC-states and mental states dictated by task. In (A), the X-axis signals time in terms of window units, and the Y-axis indicates FC-sate membership. Snapshots of windowed connectivity are represented as black dots. Underlay colors provide a visual reference for the task that subjects were performing during each window (grey = rest, blue = memo, yellow = video, green = math, white = more than one task). (D-F) show equivalent information for a second subject with poor agreement (ARI = 0.46) between FC-states and mental states. (G) Group-level average affinity matrix for all subjects with ARI > 0.8 (15 subjects out of 20)

**Figure 2.**
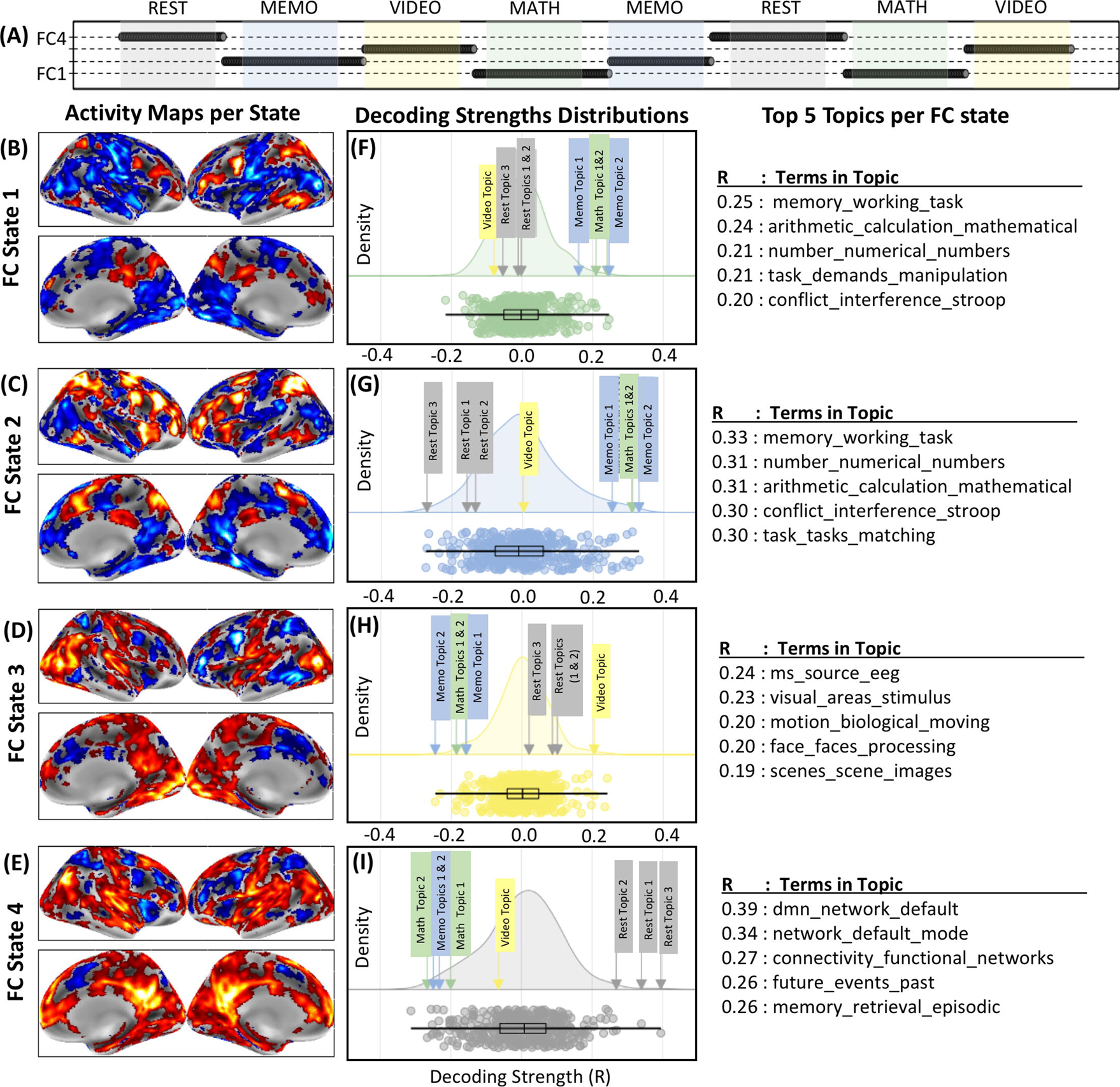
Individual subject task decoding results for the 400-topics set. (A) Scan segmentation results. Each dot represents a snapshot of windowed connectivity. The position of the dot in the y-axis indicates to which FC-state the snapshot of windowed connectivity was assigned. Tasks periods are depicted as colored bands for reference (REST: gray; MEMO: blue; MATH: green; VIDEO: yellow). (B-E) Activity maps for each FC-state obtained with SPFM. (F-I) Decoding results in the form of cloud plots and top-five lists. Cloud plots depict the probability distribution of decoding strength values across all 400 topics for each FC state in the form of kernel density estimates (colored curves), swarm plots (colored dots; one per topic) and boxplots (black). In these plots the location of “correct” topics per task are clearly marked by boxed text with arrows. Finally, the tables on the right of the figure lists the top 5 topics with the highest decoding strength for each FC state. Topic names are constructed using the top three terms associated with the topic.

Figure 1.A & D show FC timelines, in the “*all windows*” scenario, for two representative subjects with different levels of success at reproducing task timing structure. In these timelines, windows are represented as dots, with their X-axis position indicating time, and their Y-axis position indicating FC state assignments. Underlay colors provide a visual reference for the task that subjects were performing during each window (grey = rest, blue = memo, yellow = video, green = math, white = more than one task). Fig. 1.A shows a case with perfect agreement (*ARI*=1) between FC states and mental states dictated by tasks. Fig 1.D shows a case where recovery was poor (*ARI*=0.46) and FC states extended across contiguous tasks independently of their distinct cognitive demands.

Next, 3D visualizations of windowed connectivity for each subject (Fig. 1.B & E) were generated via Laplacian Embeddings (37, 38). In these visualizations, each snapshot of windowed connectivity is represented as a color-coded point in 3D space. The color of a snapshot informs us about the task, or tasks, occurring during its snapshot (grey = rest; green = math; blue = memo; yellow = visual attention; white = more than one task). The location of snapshots in these embeddings is solely a consequence of their pair-wise similarity. Task information is only used for coloring purposes, to aid with visualization, but does not contribute to the computation of the embeddings. For subjects whose FC states faithfully reproduced task states (ARI > 0.8), we observe that task homogeneous snapshots cluster together into “spoke” formations (red arrows)—one “spoke” per task—that extent away from the center of the embedding; while transition-snapshots (white dots; those spanning across tasks) form links between those spokes that go through the center of the embedding. This type of structure is less well-defined in instances of poor agreement between FC states and tasks (Fig. 1.E). Fig. 1.C & F show affinity matrices for the embeddings depicted in Fig. 1.B & E. In these matrices, a given cell represents the Euclidean similarity (*1 / [1 + Euclidean distance]*) between the location for two snapshots in 3D space. Fig 1.C shows a matrix with clear blocks of high affinity for task homogenous snapshots, in agreement with its originating embedding (Fig 1.B). This pattern becomes even more apparent when we average affinity matrices for all 15 subjects with ARI > 0.8 (Fig. 1.G).

### Data-driven estimation of number of FC States

One limitation of FC states as a model for dFC stems from the use of unsupervised clustering (most commonly *k*-*means* (34)) for their estimation, which often requires the experimenter to provide an a-priori estimation for the number of FC states (e.g., how many clusters should *k*-*means* generate). Although data-driven techniques, such as the elbow criterion, can be used to estimate this hyper-parameter; their effectiveness is limited when applied to fMRI data (22). Given how well the affinity matrices for the 3D embeddings presented in Fig. 1 represent the task structure of the multi-task scans, we decided to apply a Louvain community detection algorithm (39) to each subject matrix to estimate the number of separate communities (e.g., tasks or FC states) present in the data. For 17 out of 20 subjects, the algorithm predicted 4 communities (in agreement with the number of distinct tasks). For the remaining 3 subjects (one of them being the one depicted in Fig 1.D-F) the algorithm estimated the presence of 3 communities.

### Cognitive Correlates of FC States in the Multi-task Scenario

Cognitive decoding of FC states proceeded in two steps: 1) computation of a representative “activity” map for each FC state and 2) mapping from “activity” map into ranked lists of cognitive processes. This two-steps process was necessary for circumventing the lack of systems able to directly map FC matrices into cognitive processes.

For the computation of representative “activity” maps we relied on hemodynamic deconvolution because it does not require any information about the timing or nature of what subjects are doing in order to estimate activity—such information will not be available later when we work on the pure rest dataset. Particularly, we first used *Sparse Free Paradigm Mapping* (SPFM; (30)) to obtain voxel-wise traces of prominent activity-inducing events leading to canonical hemodynamic responses in pre-processed timeseries (red stems in Supp. Fig. 1.C & D). Next, the representative “*activity*” map for a given FC state was generated by averaging the magnitude of all activity-inducing events present within the temporal confines of that FC state on a voxel-by-voxel basis (Supp. Fig 1.D).

For the final mapping step, we relied on the reverse inference, topic-level capabilities of *Neurosynth* (31). Topics are collections of terms that appear in unison in the neuroscientific literature, and as such they provide a more meaningful level of decoding than individual terms. Decoding was attempted against two topic sets: one consisting of 50 topics and one of 400 topics (40). For each of these sets, we identified topics with a clear relationship to the tasks in the multi-task dataset. For example, TOPIC 022 in the 50 topic-set includes the terms “*memory*”, “*working memory*” and “*wm*”; which is tightly related to the 2-back (MEMO) task. Supplementary tables 1 and 2 lists all topics with a clear relationship to our tasks of interest for the 50 and 400 topic sets respectively. In addition, supplementary figures 3 and 4 depict reverse inference *NeuroSynth* activity maps for these topics.

**Figure 3.**
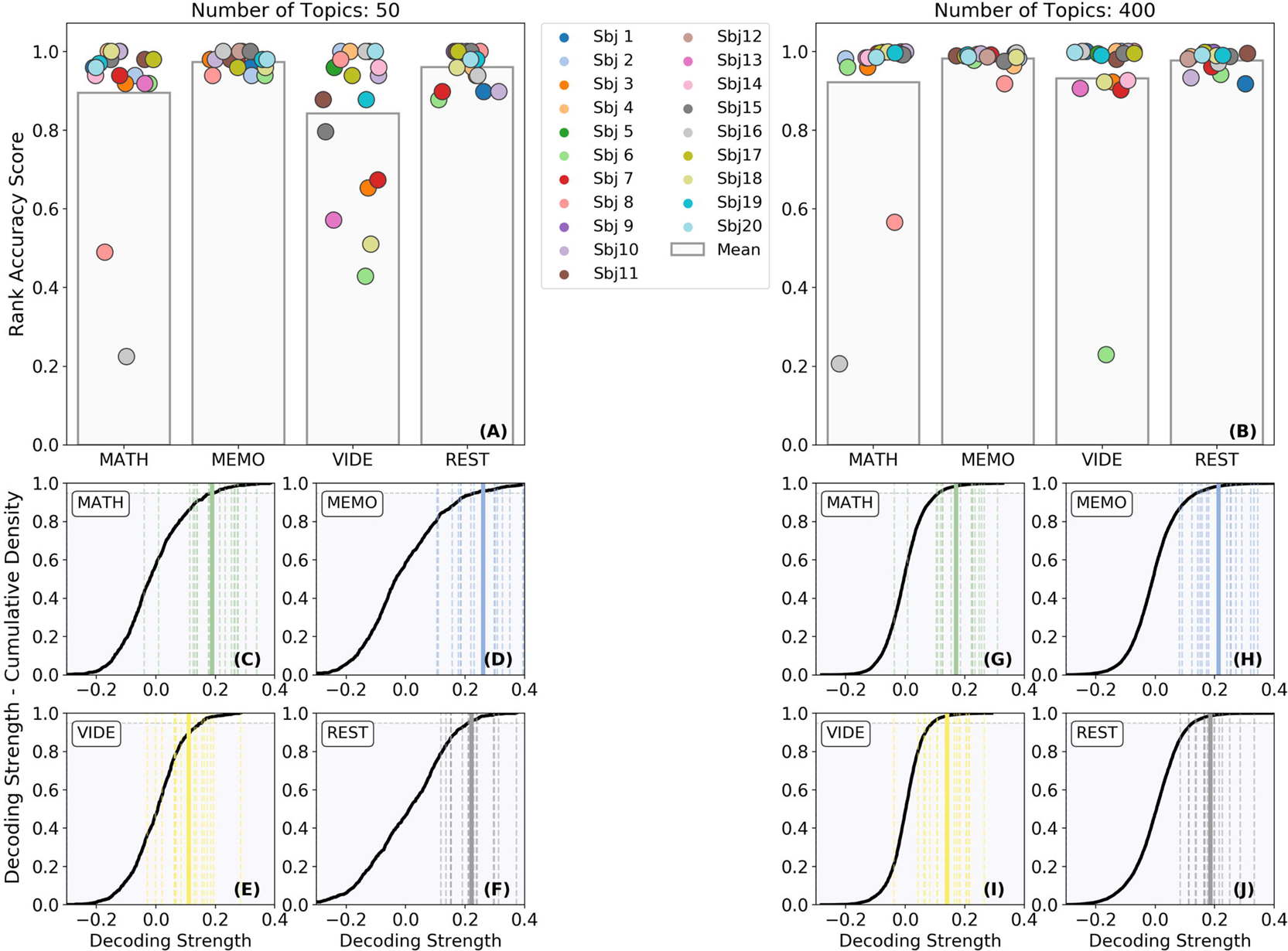
Group Evaluation of Task Decoding. (A) Rank accuracy for all subjects and tasks when using the 50-topics set. (B) Same as (A) but when using the 400-topics set. (C-F) Cumulative distribution of decoding strength values for the 50-topics set. Each panel shows results for a different task. The mean cumulative distribution across all subjects is depicted in black. Individual decoding strengths for the correct term are shown as dashed lines and the mean decoding strength across all subjects as a continuous line. In all panels, we mark the region that corresponds to 95% of the cumulative distribution with grey. (G-J) Same as (C-F) but for the 400-topics set.

**Figure 4.**
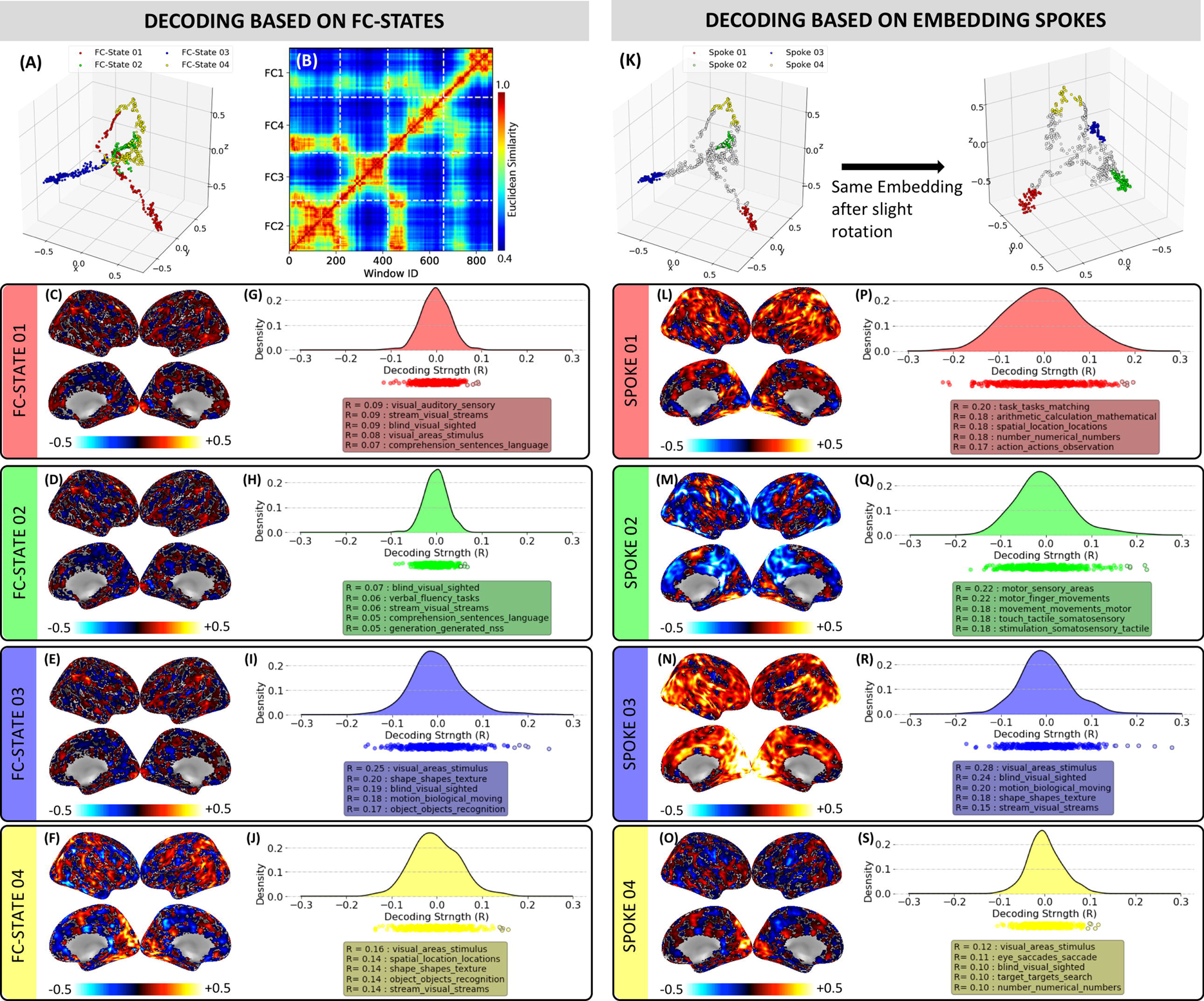
Cognitive decoding for one representative subject from the rest only dataset. (A) 3D embedding with connectivity snapshots colored according to automatically extracted FC states. (B) Affinity matrix for the embeddings in panel A. (C-F) Activity maps computed via SPFM for each of the FC states. (G-J) Distributions of decoding strengths for the 400-topics set and lists with the top-5 topics for each FC-state. (K) Same embedding as in (A), but this time colors signal the windows assigned to spoke-like structures following the procedures described in the SI materials. Two different views (different rotation angles) are presented to help better appreciate the structure of the embedding. (L-O) Activity maps computed using only the windows identified as members of spoke-like structures (e.g., non-white windows). (P-S) Distribution of decoding strengths for the maps shown in L-O.

Fig. 2 and Suppl. Fig. 5 show decoding results for two representative subjects using the 400-topics set (Fig. 2) and the 50-topics set (Supp. Figure 5). Both figures have the same structure. Panel A shows FC state timelines. Panels B-E shows “*activity*” maps for each FC state. Panels F-I show decoding results per FC state both as a table listing the top-five topics with the highest decoding strength and as cloud plots with the distribution of decoding strength across all available topics. For qualitative evaluation purposes, the location of correct topics (i.e., those with the most direct relationship to the task; see Suppl. Tables 1 & 2) are clearly marked for all FC states in panels F-I. In particular, for *Sbj17* (Fig. 2), we can observe how the FC state timeline faithfully follows the experimental task timing (*ARI*=1.0). For this subject, FC state 1 spanned periods of MATH and FC state 2 periods of the MEMO task. Activity maps for these two states contain prominent clusters of activity in dorsolateral pre-frontal cortex and parietal regions (Fig. 2.B & C); consistent with tasks with a heavy working memory component. Similarly, FC state 3, which overlaps in time with the visual attention task (VIDEO), has prominent areas of activity around the MT/V5 region and other visual regions in ventral temporo-occipital cortex (Fig. 2.D). Finally, for FC state 4—which spans rest periods—we observe activity in regions of the default mode network (Fig. 2.E). As for the actual decoding, for FC states 1 and 2, we observe topics with terms such as “*memory*”, “*working*”, “*arithmetic*”, “*calculation*” and “*numbers*” among the top decoding topics (Fig 2.F & G). For FC state 3 (VIDEO), top topics include the terms “*visual*”, “*motion*”, “*biological*”, “*moving*” and “*scenes*” (Fig. 2.H). Finally, for FC state 4 (REST), the three topics that best describe rest (see Suppl. Table 2) appear as the top three decoding terms for this FC state (Fig 2.I). Similar results can be seen in Suppl. Fig. 5 when decoding was done using the 50-topics set on a different representative subject.

**Figure 5.**
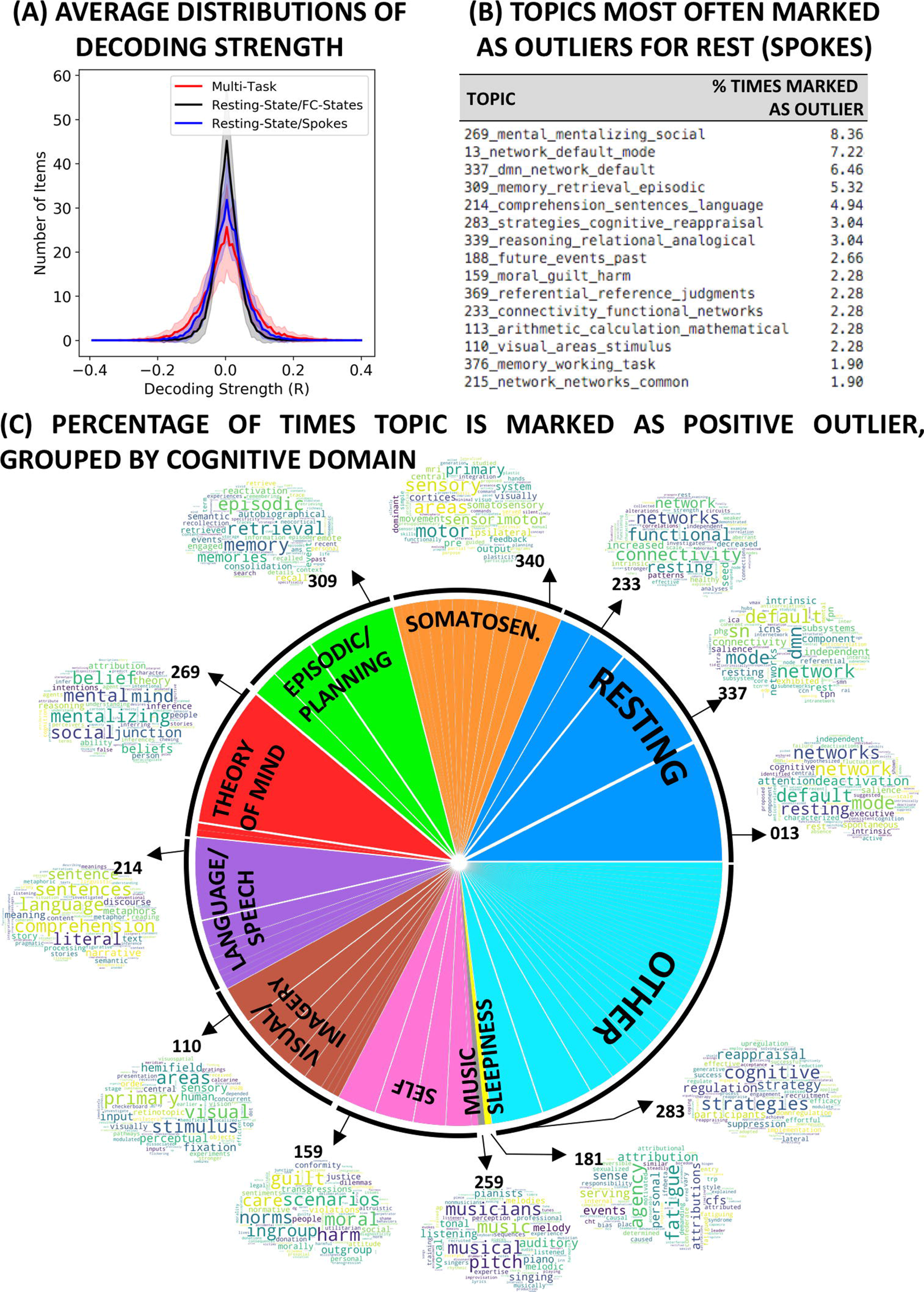
(A) Average decoding strength distributions for the 400 topic-set under three different scenarios: (red) FC-states detected in the multi-task dataset; (black) FC-states detected in the pure rest scenario; (blue) windows in the vicinity of spoke-like structures in the pure rest scenario. (B) List of the top 15 topics marked as positive outliers (e.g., signaled to have a strong relationship with provided activity maps) during the decoding of spoke-like structures in pure rest 3D embeddings. Topic names were constructed concatenating the topic ID in NeuroSynth with the three top terms part of the topic. The percentage on the right indicate the number of times the topic became an outlier relative to the total number of outlier detection in the whole sample. (C) Depiction of the distribution of outlier topics for spoke-like structures in the pure rest scenario grouped by cognitive domains previously reported to describe most common cognitive processes subjects engage with during rest.

Fig. 3 shows group-level task decoding results. Fig. 3.A-B show rank accuracy (41)—a measure of how often the correct topics appeared at the top of the decoding rank—for all task segments and subjects as colored dots, and across-subject mean rank accuracy per task as grey bars for the 50-topics set (Fig. 3.A) and the 400-topics set (Fig. 3.B). Mean rank accuracy was above 0.8 for all tasks for most subjects. Worse decoding results were obtained for the *visual attention* (VIDEO) task when using 50-topics. Overall, decoding was more successful when using the 400-topics set relative to the 50-topics set; this despite a large increase in the ratio of wrong-to-correct choices. In all instances, mean rank accuracy was significantly higher than chance (*p*<0.05; see SI for details). Figure 3.C-J show average cumulative distributions of decoding strength for the different tasks (black curves). In addition, individual decoding strength for correct topics are depicted as dashed colored lines. A colored continuous bold line marks the mean correct decoding strength across all subjects. In all instances, except for VIDEO using the 50-topics set (panel 3.E), the mean decoding strength for correct topics fell to the right of 95% of the cumulative distribution of decoding strengths (shaded region).

### Periods of Distinct Cognition during Rest

Next, we applied the same methods to a subset of twenty 15 mins-long eyes-open resting scans acquired on a different 7T system part of the HCP (32). Subjects were selected to minimize motion and sleepiness confounds (detailed selection criteria in SI materials). Suppl. Fig. 6 show traces of windowed eye closure times per subject. It can be observed that all subjects remained awake for the whole scan duration. Supp. Fig 7 shows traces of absolute and relative motion for all 20 subjects. Average absolute motion was 0.27 ± 0.18 mm, and average relative volume-to-volume motion was 0.10 ± 0.07 mm.

Figure 4 summarizes results of these analyses for one representative resting subject (*sbj05*). Panel A shows the 3D embedding for this subject, with connectivity snapshots colored by FC state (3D embeddings for all rest subjects in Suppl. Figure 8.A). Number of FC-states for the rest datasets varied between three and four. Large inter-subject variability in 3D embedding patterns can be observed in Suppl. Fig. 8.A for this second dataset. Although “*spoke-like*” structures are present for some subjects (e.g., the one in Fig. 4), this motif is not as prominent as in the multi-task dataset (Supp. Fig. 8.B). Moreover, FC-states in rest scans do not necessarily align with distinct spokes in the same manner they did in the multi-task scans (e.g., in Fig. 4.A, FC1 extends into a second spoke that is shared between FC1 and FC4). Panel 4.B shows the affinity matrix for the 3D embedding shown in 4.A. Dashed lines identify the confines of estimated FC-states, which were very similar in size and contain only temporally contiguous windows. This behavior was common across the majority of rest scans, suggesting that FC-states did not fully capture the structure depicted by the corresponding 3D embeddings.

Panels 4.C-F show “*activity*” maps for all four FC-states detected in *sbj05*. Activity maps are very similar across states, particularly for FC-states one, two and three. Panels 4.G-J show corresponding distributions of decoding strength per FC-state for the 400-topics set. Distributions are often narrower than in the multi-task scenario, suggesting fewer topics show strong positive associations with activity maps. This is particularly clear for FC states one and two, whose top-ranking terms have decoding strengths below 0.1. This same narrowing effect can be observed at the group level by looking at group-averaged distributions of decoding strength (Fig. 5.A). Average distributions are wider and have longer tails for the multi-task dataset (red) than for the rest dataset (black). We quantified this effect by looking at the number of topics that constitute positive outliers for each FC-state (i.e., those above the third quantile plus 1.5 times the interquartile range) in both scenarios. We found that there are significantly more outlier topics for the multi-task dataset than the rest dataset (*T=2.74; p=0.007*).

Lastly, to explore if observable spoke-like structures in pure rest embeddings correspond to periods of distinct cognition—even if automatically estimated FC states were not able to effectively capture them—we decided to apply our cognitive state decoding method focusing only on windows sitting at the distal ends of rest 3D embeddings. For each subject, we found the *x* most distal windows in the embedding (in terms of Euclidean distance), with *x* being equal to the number of FC-states. Next, for each distal end, we selected all windows sitting within a given distance *d* from it (*d* = average inter-snapshot distance for all task-homogenous windows in the multi-task dataset). This procedure allowed us to automatically identify sets of spatially contiguous snapshots sitting near the distal ends of the embeddings. Fig 4.K shows the output of this procedure for *sbj05*. Both 3D plots in the panel show the same data viewed from different angles. Suppl. Fig. 9 shows the same information for all rest subjects. Next, we took these newly defined groups of windows and proceeded with the cognitive decoding step. Fig. 4.L-O show activity maps for the four groups of windows sitting at distal ends of the embedding for *sbj05*. Activity maps are now more clearly differentiable across clusters. The map for *Spoke01* shows activity in lateral parietal and frontal regions. The map for *Spoke02* has activity mostly focused on sensory motor regions. The map for *Spoke03* presents strong activity in occipital, limbic and insular regions. And, finally, the map for *Spoke04* shows activity primarily concentrated in the calcarine region. Fig 4.P-S shows corresponding decoding results. Distributions now have widened and have longer tails, suggesting activity maps are now more strongly associated with distinct topics. This effect is also appreciable at the group level (Fig 5.A – blue trace). Consequently, decoding strengths for top topics are larger; with all top 5-topics having R > 0.1. Moreover, top topics are now associated with distinct cognitive domains. For example. *Spoke01* contains topic associated with numeric mental activity (e.g., “*arithmetic_calculation_mathematical*” and “*number_numerical_numbers*”). *Spoke02* seems to encompass periods of strong somatosensory activity, as indicated by the terms “*motor_sensory_areas*”, “*motor_finger_movements*”, “*movement_movements_motor*”, etc. Finally, *Spoke03* and *Spoke04* point to scan segments dominated by strong visual activation perhaps associated with visual imagery.

Across all subjects, out of the 400 available topics, only 78 topics became positive outliers for one or more spokes. Fig 5.B lists the top 15 topics most commonly marked as outliers. This includes topics clearly associated with pure rest (topics 12, 337, 233, 215), thinking about the past or future (topics 309 and 188), theory of mind (topic 269), language/inner speech (topic 214), self-evaluation (topics 159 and 369), mental computations (topics 113 and 376), and visual activity (topic 110). Those all correspond to cognitive domains previously reported as describing the types of mental processes subjects commonly engage with during rest (26, 42, 43). Finally, Fig 5.C provides a summary view of all topics marked as positive outliers for spoke-like structures in pure rest scans. Topics are grouped into the above-mentioned cognitive domains describing resting mental activities. Approximately 75% of topics fall within one of these categories: resting (17.87%), somatosensory (9.89%), episodic/planning (9.51%), theory of mind (9.13%), language/speech (9.13%), visual/imagery (9.13%), self (8.37%), music (0.4%) and sleepiness (0.4%).

## DISCUSSION

Resting dFC is an empirically observed phenomenon with promising translational value (4, 5) for which several methodological and mechanistic questions remain unresolved (24). We address some of them here. First, we show evidence in support of the hypothesis that on-going self-paced cognition is a contributor to dFC phenomena during rest, although the degree of contribution may vary across subjects, as well as across cognitive domains. Second, we demonstrate that data-driven estimates of FC states based on fixed-sized sliding window approaches—one of the most prominent modeling approaches for dFC—do not always capture periods of distinct cognitive processing during rest, even though FC states robustly align with cognitive processes imposed by external task demands (i.e., multi-task scenario). Overall, our results highlight the cognitive relevance of resting dFC, how a single FC static structure may not be sufficient to capture the diversity of cognitive processes occurring during rest, and the need to update the definition, computation and thus interpretation of FC states, especially when it comes to how they may relate to cognitive processing during rest.

### Cognition contributes to observable dFC during Rest

Snapshots of time-varying FC spanning task-homogenous periods in the multi-task scenario clustered together at the distal ends of spoke-like structures in the 3D embeddings (Fig. 1; Suppl. Fig. 8.B). This motif of similar elements sitting together at corners of Laplacian embeddings has been previously reported for different domains, including speech (37) and neuronal spike sorting (44). For the HCP rest scans, similar structures could be observed in a subset of subjects, suggesting the presence of isolated periods of homogenous FC distinct from other portions of the scan. Moreover, reverse-inference results suggest that such periods are associated with different cognitive processes (Figs. 4 & 5). Overall, these results support our working hypothesis that covert on-going cognition affects short term estimates of FC in a similar manner to how overt externally imposed cognition does (3); and, subsequently, that cognitively meaningful dynamics can be a contributing factor to resting dFC.

In the multi-task scenario, we were able to objectively validate cognitive inferences. Unfortunately, our ability to do so for rest scans is severely limited. Nonetheless, the cognitive processes inferred for spoke-like structures in rest embeddings agree with previous reports of the most common mental operations subjects undertake during rest (26, 42, 43), and different spokes tend to be associated with a different set of mental processes (Fig. 4). Future work could better validate the neural accuracy of the cognitive mappings. For example, concurrent skin conductance, electromyography and video recordings of subjects in the scanner bore may help validate inferences regarding periods of somatosensory activity, self-generated body movements (29) or strong emotional content. Additional validation may be possible using rest scans accompanied by retrospective descriptions of subject’s mental processes during the scan. Delamillieure et al. (26) suggested that subjects can be grouped according to the most prominent type of mental activity they conduct while resting. Future work should evaluate if similar groupings can be obtained on the basis of retrospective reports and data-driven decoding. Finally, dFC phenomena of limited complexity have been reported in the absence of consciousness (16, 45). Should spoke-like structures in dFC embeddings be the manifestation of covert cognitive processes, those structures should dissolve for rest scans conducted under deep anesthesia.

The timing and nature of cognition occurring during rest is heterogenous across subjects (26, 43). The same is true for other postulated contributing factors such as arousal, motion or sleep. The embeddings presented here demonstrate that resting dFC is quite heterogenous across subjects even when both motion and sleep contributions were minimized via sample selection. Future work should help isolate all these factors and quantify their relative contributions at the individual subject level. For example, templates of regional activity correlated with pupil dilation may help isolate the contributions of arousal (46). Such quantification efforts are key for merging current discrepant views on the origin of rest dFC. As previously mentioned, Laumann et al. (25) recently stated that rest dFC is primarily the result of sampling variability, arousal fluctuations, sleep and motion. One supporting argument made by the authors was the observation of limited changes in multi-variate kurtosis for pure rest scans as compared to task/rest alternating scans (Fig. 6 in (25)). Yet, reported histograms of multi-variate kurtosis for pure rest scans (blue) have a long right tail and bump that overlaps with the kurtosis distribution for alternating task/rest scans (green and red traces). Our results suggest that cognition may contribute to resting dFC differently across subjects, with potentially negligible contribution in some subjects (those with very diffuse spokes). It could be argued that the right tail of the distribution of multi-variate kurtosis values reported in (25) for pure rest (Fig 6, blue trace) represents the presence of a limited group of subjects with strong contributing on-going cognition. Moreover, assessing the contribution of different factors will not only help settle this argument, but also understand the origin of previously reported systematic differences in dFC across normal and patient populations (2, 18, 19). For example, it may be the case that differences in both static or dynamic estimates of FC across populations are not solely indicative of differences in intrinsic patterns of connectivity but may be also contributed by subjects from different populations engaging systematically into different forms of cognitive processing as they lay inside the scanner. For example, depressed patients may engage more often in regurgitative thinking; while a different population may be more prone to perform mathematical computations as a form of distraction while being scanned.

### Open-ended Cognitive Inference for fMRI Scans

Here we describe and test a pipeline for blindly inferring periods of distinct cognition occurring during fMRI scans. The primary components of the pipeline are: 1) dFC estimates are used to temporally segment scans into FC homogenous periods; 2) hemodynamic deconvolution is used to generate activity-like maps for each period; and 3) activity-based reverse inference is used to deduce the cognitive processes occurring during those periods. All these steps can be performed on individual scans and require no training dataset. As such, decoding is not limited to group-level inferences and the breadth of cognitive states to be inferred is ultimately set by the breadth of the neuroimaging literature mined by the *NeuroSynth* platform.

Nonetheless, each of these steps set limitations regarding the specificity and sensitivity of resulting inferences. For example, the temporal segmentation step is designed to accommodate more than one cognitive state per scan. Unfortunately, the present results suggest its ability to effectively do so in a meaningful manner is limited to externally imposed cognition. Similarly, the correctness of the hemodynamic deconvolution step will depend on the degree of correspondence between canonical and actual hemodynamic response shapes (which will vary per location and subject). Finally, limitations inherent to the *NeuroSynth* platform—such as limited specificity of topics, vocabulary biases (31, 47) also apply here. Future developments should help address many of these limitations. For example, multi-echo fMRI can improve the accuracy of the deconvolution (48); and improved probabilistic decoding frameworks may increase the specificity of the inferences (47). In addition, it will be valuable to develop systems able to directly map FC matrices into cognitive states in an open-ended fashion, erasing the need to generate representative “activity” maps for each FC state as an intermediate step. The development of such systems is intricate given the diversity of brain parcellations used to compute functional connectivity matrices and the fact that matrices tend to be reported only in graphic form—i.e., images—and not as numerical arrays. Developments in image recognition that could automatically mine and transform images of FC matrices into numerical arrays, and novel meta-analytic tools able to merge connectivity information from diverse parcellations may make those systems readily available in the near future.

### FC states as a Model for Resting dFC

FC states currently constitute a mainstream approach to model, summarize, and report within-scan dFC. Repeatedly, researchers have reported on the behavioral (3, 49) and clinical relevance (2, 50) of this model. Yet, FC states present important limitations resulting from the use of fixed-size sliding windows and *k*-*means* as key steps in their estimation. First, the FC-states framework commonly assumes that dFC is effectively characterized in terms of a limited set of distinct FC configurations (those detected by *k*-*means*). As the multi-task embeddings demonstrate (Fig 1), even if that were true, the limited temporal resolution of sliding windows will often result in adulterated FC estimates—spanning several unique cognitive configurations—that sit at interspaces between clusters (white dots) and diffuse true clustering structure in the data (task-colored dots). Because *k*-*means* enforces membership (all inputs must be assigned to a cluster) and tends to generate clusters of equal size when true cluster structure is diffused or missing, k-means is not effective at capturing periods of distinct FC in the rest scenario (Fig. 4.A). This is clearly exemplified by the fact that, for the multi-task scenario, the ARI significantly decreased when all overlapping windows enter the analysis relative to when only task-homogenous windows do (Suppl. Fig. 2). To avoid such issues, future FC state modeling should rely on variable size windows that align with FC transitions (51) or rely on non-windowed estimates of time-varying connectivity such as dynamic conditional correlation (52). In addition, clustering techniques than can accommodate for both multiple membership (e.g., one element can be part of two or more different clusters) and null membership (inputs may be assigned to no cluster at all) should also be adopted (53, 54). More importantly, future research should elucidate whether dFC should be conceptualized as a continuous or discrete phenomenon (55). Ultimately, it may be that a hybrid framework able to accommodate both the presence of discrete, transient meaningful configurations (signaling distinct periods of cognition or sensory activity whenever present) and underlying meta-states (potentially capturing intrinsic maintenance and exploratory processes) may be the most appropriate conceptualization (56, 57).

### Extensions to Prior Work, Limitations, Future Directions

Our original study conducted on this multi-task dataset demonstrated that FC states derived from non-overlapping task-homogenous windows robustly recovered temporally disjoint periods of homogenous cognition (3). Here we extend those results in several ways. First, we demonstrate a robust data-driven way to infer the number of states (i.e., tasks) using affinity matrices in the embedded 3D space. In our original work, the number of states was provided by the experimenters. Second, we not only describe a way to temporally segment multi-task scans into task homogenous periods, but also how to deduce the cognitive processes underlying those periods in an open-ended fashion that requires zero timing information or training data. Third, we demonstrate that those results can be accomplished even if overlapping task-inhomogeneous windows enter the analyses. In the original study, the onset/offset times of task blocks were used to restrict analyses to only task-homogenous non-overlapping windows.

Several limitations, in addition to those regarding imperfect ability for validation in the resting scenario, apply here. First, dFC phenomena most likely resides on a higher dimensional space beyond three dimensions. Three dimensions were selected for the current analyses because of their representational ease and because they were able to capture task structure much better than their 2D counterparts (Supp. Fig. 10). Yet, 3D embeddings were not able to explain all clustering errors. For example, Fig. 1.D&E shows an example of how 3D embeddings were able to capture the proximity of snapshots from the first MEMO, VIDEO and MATH blocks in the original higher dimensional space leading to their mistakenly joint membership to FC-State 1; but failed at capturing the same information regarding the mistaken joint assignment of windows from the second MATH and VIDEO blocks to FC-state 3. Higher dimensional spaces may better capture all relationships of interest as they will not distort original pair-wise relationships so abruptly. Yet, tools for visualizing such spaces are limited and may be difficult to interpret. Future work should evaluate the effectiveness of working with higher dimensional spaces, and the ability of visualization tools such as parallel coordinates (58) and glyphs (59) to render meaningful representations of dFC in such higher dimensional spaces. In addition, future studies should evaluate the ability of other manifold learning methods, such as T-SNE (60), to aid in the visualization and interpretation of dFC phenomena during task and rest. A second limitation of the current study is that we only worked with one implementation of FC-states. We previously demonstrated that methodological decisions can significantly alter the strength of the relationship between FC-states and cognitive states dictated by tasks (3). As alternative methods to estimate FC-states exist in the literature (1, 53), future work should elucidate how well our conclusion of limited ability of FC-states to capture periods of distinct cognition during rest generalizes to these other variants of FC state modeling.

Despite the above-mentioned limitations, we believe that the current results advance our understating of dFC during both rest and task. They provide evidence in support of the hypothesis that resting dFC includes manifestations of covert cognition; suggesting that several different behaviorally relevant whole-brain FC configurations may occur during a single rest scan. In addition, we confirmed that data-driven estimates about the cognitive nature of most common mental processes occurring during rest agree with those inferred via retrospective questioning in other subject samples. Finally, our results also underscore how differences in effect size and temporal characteristics of externally-imposed and self-driven cognition may obstruct direct translation of methods and conclusions across both scenarios.

## METHODS

A detailed description of the data and methods can be found in the SI materials accompanying this document.

## ACKNOWLEDGEMENTS

This research was possible thanks to the support of the National Institute of Mental Health Intramural Research Program. Portions of this study used the high-performance computational capabilities of the Biowulf Linux cluster at the National Institutes of Health, Bethesda, MD (*biowulf.nih.gov*). This study is part of NIH clinical protocol number NCT00001360 and protocol ID 93-M-0170. CCG was supported by the Spanish Ministry of Economy and Competitiveness through the Juan de la Cierva Fellowship (IJCI-2014-20821) and the “Severlo Ochoa” Programme for Centres/Units of Excellence in R& D (SEV-2015-490). Resting-state data were provided by the Human Connectome Project, WU-Minn Consortium (Principal Investigators: David Van Essen and Kamil Ugurbil; 1U54MH091657) funded by the 16 NIH Institutes and Centers that support the NIH Blueprint for Neuroscience Research; and by the McDonnell Center for Systems Neuroscience at Washington University. Finally, JGC would like to thank his parents, Enrique Gonzalez and Maria Margarita Castillo, and wife, Robyn Russo, for the love, courage and strength demonstrated during the toughest times.

## Footnotes

1 This multi-task dataset consists of 25 mins long scans acquired as subjects engage and transition between four different tasks—namely rest, math, 2-back (MEMO) and visual attention (VIDEO)—distributed in 3 mins long periods (two such periods per task).

2 The ARI ranges from 1 to below 0, with 1 indicating perfect recovery of clusters. Ranges established in the literature describe an *ARI* > 0.9 as excellent, 0.9 > *ARI* > 0.8 as good, 0.8 > *ARI* > 0.65 as moderate, and ARI < 0.65 as poor recovery, respectively 36. Steinley D (2004) Properties of the Hubert-Arabie adjusted Rand index. *Psychol Methods* 9(3):386-396.

