## Supplementary Materials for "Imaging the spontaneous flow of thought: Distinct periods of cognition contribute to dynamic functional connectivity during rest"

### METHODS

#### Multi-Task Dataset

The multi-task dataset used here has been previously described in detail in (1). In summary, it contains data from 22 subjects (13 females; age 27 +/- 5 y.o.) who gave informed consent in compliance with a protocol approved by the Institutional Review Board of the National Institute of Mental Health in Bethesda, MD. The data from two subjects were discarded from the analysis due to excessive spatial distortions in the functional time series.

The MRI data have been deposited in Xnat Central, https://central.xnat.org (project ID: FCStateClassif).

#### Multi-Task Experimental Paradigm

Subjects were scanned continuously for 25 min and 24 s while performing four different tasks: rest with eyes open (*REST*), simple mathematical computations (*MATH*), 2-back working memory (*MEMO*), and visual attention/recognition (*VIDEO*). Each task occupied two separate 180-s blocks, preceded by a 12-s instruction period. Task blocks were arranged so that each task was always preceded and followed by a different task. Additional details can be found on the supplementary materials accompanying Gonzalez-Castillo et al. (1).

#### Multi-Task Data Acquisition

Imaging was performed on a Siemens 7 Tesla MRI scanner equipped with a 32-element receive coil (Nova Medical). Functional runs were obtained using a gradient recalled, single shot, echo planar imaging (gre-EPI) sequence: (TR=1.5 s; TE=25 ms; FA=50°; 36 interleaved slices; slice thickness = 2 mm; in-plane resolution = 2×2 mm; GRAPPA=2). Each multi-task scan consists of 1,017 volumes acquired continuously as subjects engage and transition between the different tasks. In addition, high resolution (1mm$^3$) T1-weighted magnetization-prepared rapid gradient-echo and proton density (PD) sequences were acquired for presentation and alignment purposes.

#### Multi-Task Data Preprocessing

Data preprocessing was conducted with AFNI (2). Preprocessing steps match those described in Gonzalez-Castillo et al. (1), and include: despiking; physiological noise correction (in all but four subjects); slice time correction; head motion correction; regression of additional nuisance signals according to the ANATICOR model (3); bandpass filtering [0.03 – 0.18Hz]; and spatial smoothing (*FWHM* = 4mm). The cutoff frequency of the high pass filter was chosen to match the inverse of window length (*WL* = 30s); following recommendations from Leonardi and Van De Ville (4).

In addition, spatial transformation matrices to go from EPI native space to Montreal Neurological Institute (*MNI*) space were computed for all subjects following procedures previously described in (5). These matrices were then used to bring publicly available regions of interest (ROI) definitions from MNI space into each subject's EPI native space.

#### Resting-State Dataset

In addition to the previously described multi-task scan dataset, we also conducted analyses on a set of 20 resting-state scans part of Human Connectome Project (6). In particular, we used the first resting-state scan [TR=1s; TE=22.2ms; FA=45°; Voxel Resolution = 1.6x1.6x1.6mm$^3$; Multiband Factor = 5; GRAPPA = 2] from 20 subjects part of the 1200 Release (March 2018) for which FIX (7) pre-processed resting-state scans acquired on a 7T MR system were available. Subject selection proceeded as follows. First, out of the 175 subjects available, we selected the 144 subjects that completed both the NIH and non-NIH toolbox behavioral batteries, as well as alcohol and smoking questionnaire, and the family history of psychiatric and neurological disorders. Of these remaining 144 subjects, we further restricted our sample to the subset of 106 subjects for which eye tracking data was available for the first resting scan. Next, based on windowed estimates of percent time of eye closure per window, we selected the 37 subject subjects that never had their eyes closed for more than 40% of the duration of any sliding window utilized in our analyses (Suppl. Fig. 6). Finally, we restricted our study to the 20 subjects with the least amount of maximum absolute motion among those 37 subjects (Suppl. Fig. 7).

In addition to the pre-processing performed via the HCP FIX-ICA denoising pipeline, we also applied band-pass filtering (0.03 – 0.18Hz]), and spatial smoothing (*FWHM* = 4mm) to match equivalent filtering and spatial smoothing steps performed on the multi-task dataset. Following these steps, scan segmentation analyses proceeded as described below for both datasets (e.g., multi-task and rest).

#### Scan Segmentation into Cognitively Homogenous Segments

Segmentation of multi-task scans into cognitively homogenous segments was performed on the basis of short-term 30 s long functional connectivity patterns (snapshots) following procedures previously described in Gonzalez-Castillo et al. ((1); Supp. Fig. 1.B). First, representative time-series ROIs from the Craddock (8) 200-ROI atlas were obtained and input to a PCA (keep 97.5% of the variance) to reduce the dimensionality of the data. This resulted, on average, in connectivity matrices of size 72 x 72 instead of the original 200 x 200. Remaining PCA time series were subsequently segmented in time using overlapping windows of duration 30s. Sliding step was set to 1.5 s (one *TR*). For each window, we computed all pairwise correlations between PCA time series, transform them into Z-scores using the Fisher transform, and put them in vector form. For each multi-task scan this procedure resulted in 988 vectors (one per-window). We often refer to these vectors of windowed connectivity as connectivity snapshots or simply snapshots throughout the manuscript.

Lastly, these connectivity vectors were inputted into the *k-means* (*k*=4 given the number of tasks; distance metric=correlation) clustering algorithm in MATLAB 2017b, which groups connectivity vectors into four groups by maximizing within-group similarity and between-group dissimilarity. We hypothesize that each group will contain primarily vectors from the same task based on the original results from Gonzalez-Castillo et al. (1). If so, the time spanned by all windows in a given group define a segment from the original scan during which the subject was engaged with primarily one type of cognitive activity (e.g., one task). As an unsupervised clustering algorithm, k-means only informs us about groupings in the data, not about the meaning (e.g., the task or main cognitive process) associated with each group.

#### Computation of Activity Map per Segment

Pre-processed time series—excluding the low pass filtering step—were also input to AFNI program *3dPFM*, which implements the fMRI deconvolution algorithm named Sparse Free Paradigm Mapping (SPFM; (9)). This program takes as input voxel-wise fMRI time series and a canonical hemodynamic response model—here we used the SPFM canonical response (SPGM1 function on AFNI program *3dDeconvolve*). The output consists on voxel-wise time series of sparse activity-inducing events (Suppl. Fig. 1.B) that best explain the input fMRI time series. The deconvolution algorithm enforces sparsity in time via an $L_1$ norm regularized estimator using the LASSO. Here, the Akaike's Information Criterion (*AIC*) was used to select the regularization parameters per voxel.

Voxel-wise time-series of activity inducing events were computed with AFNI program *3dPFM* using entire scan time series. Subsequently, activity-like maps were computed for each cognitively homogenous scan segment as dictated by the k-means analysis described above. In these maps, the intensity of a given voxel corresponds to the average intensity of all activity-inducing events detected by *SPFM* that fall within the temporal constrains of the given segment (see Suppl. Fig. 1.D). A such, this procedure resulted on one "activity" map per cognitively homogenous segment. These maps constitute the input to the decoding step described next.

#### Cognitive Decoding

To minimize limitations in terms of the number of cognitive processes that could be decoded, we decided to rely on existing meta-analytical databases of brain activity for this last step of the analysis. More particularly, we decided to use the reverse inference capabilities of the *Neurosynth* system; which permits decoding at two different levels: that of individual terms (10) or that of topics (11). Because topics are collections of terms that frequently co-occur in the neuroscientific literature, they provide a more meaningful level of decoding; and were chosen as the decoding target here. Two different topic sets were used: the 50- and 400-cognitive state topic sets described in (11). In this manner, we were able to evaluate the efficiency of the method under two different difficulty levels.

Topic-level decoding with *Neurosynth* proceeds as follows. First, for each topic set, *Neurosynth* computes a reverse inference map (see Suppl. Figs. 3 & 4 for some examples) based on activity foci reported in large corpus of neuroscientific literature as described in (11, 12). Next, when an instance for decoding (e.g., the maps generated in "Computation of Activity Map per Segment") is available, *Neurosynth* computes the spatial correlation between this input and all available topic maps. This correlation informs about the likelihood of a given topic being associated with the input map. The output of *Neurosynth* usually consists on a sorted list of topics and their associated correlations with the input. The higher the "correct" topic for a given input appears in this rank, the more accurate the decoding was. Below we describe how we evaluated the quality of the decoding.

#### Evaluation of Scan Segmentation and Decoding

##### *Scan Segmentation Evaluation*

To quantitatively evaluate success at recovering the periods during which subjects were performing the same mental task, we used the adjusted rand index (*ARI*) (13). The *ARI* measures quality of data-driven clustering (k-means) against an existing gold-standard (experimental tasks). It ranges from 1 to below 0, with 1 indicating perfect recovery of clusters. Ranges established in the literature describe an *ARI* > 0.9 as excellent, 0.9 > *ARI* > 0.8 as good, 0.8 > *ARI* > 0.65 as moderate, and *ARI* < 0.65 as poor recovery, respectively (14).

Clustering was attempted under two scenarios: (a) considering all available windows; and (b) after removal of all non-task homogenous windows (e.g., those spanning one or two tasks plus instruction periods). Independently of the scenario, the *ARI* was computed only considering task-homogenous windows. In other words, no matter which windows entered the k-means analysis, the *ARI* was computed only based on the grouping of task-homogenous windows as those are the only ones for which an unambiguous correct answer clearly exists.

##### *Decoding Evaluation*

Decoding was performed using two different topic sets: 50 and 400 cognitive topic sets. These two sets contain one or more topics that strongly relate to the four tasks under scrutiny. For example, within the 50 topic-set, only one topic (*TOPIC 022*) contains the terms "*memory*", "*working memory*", "*wm*". Accordingly, decoding success for the *MEMO* segments is quantified in terms of the decoding ranking for *TOPIC 022*. Similarly, within the 400 topic-set, two topics (*TOPIC14* and *TOPIC376*) contains those terms. In this case, decoding success for *MEMO* segments will be quantified considering the average decoding ranking for these two terms. Suppl. Tables 1.A and 1.B list the "correct" topics for our four tasks in the 50 and 400 topic-set, respectively. *Neurosynth* reverse inference maps for all "correct" topics are displayed on Supplementary Figures 3 (50-topic set) and 4 (400-topic set).

Supplementary Table 1: Correct topics for all tasks in the 50 topic-set

| Task | Topic | Top Terms | |
|---|---|---|---|
| **Math** | 042 | 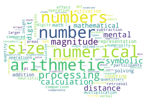 | number, numerical, arithmetic, numbers, size, processing, magnitude, calculation, symbolic, mental, distance, mathematical, digit, representation, solving, activation, math, counting, addition, multiplication, retrieval, operations, comparison, effect, set, participants, subtraction, operation, numerosity, estimation, strategies, heuristic, compared, areas, quantity, digits, quantifiers, solution, insight, larger |
| **Memo** | 022 | 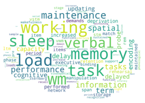 | memory, working, wm, load, task, sleep, verbal, maintenance, performance, information, spatial, term, tasks, cognitive, delay, encoding, capacity, probe, delayed, updating, deprivation, executive, performed, increased, manipulation, storage, stm, vstm, items, span, retention, item, rehearsal, phase, match, demands, rem, period, maintained, vwm |
| **Video** | 011 | 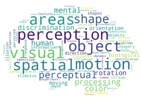 | motion, visual, perception, areas, spatial, object, shape, perceptual, processing, color, mental, human, discrimination, rotation, biological, orientation, moving, information, imagery, stimulus, objects, direction, responses, shapes, stream, illusion, participants, sensitive, form, static, texture, depth, perceived, response, location, experiment, colour, dynamic, dimensional, selective |

| | | | |
|---|---|---|---|
| **Rest** | 010 | 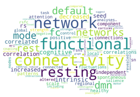 | connectivity, functional, network, resting, networks, default, dmn, mode, rest, fc, intrinsic, cognitive, correlation, increased, correlated, seed, rs, decreased, independent, correlations, spontaneous, connections, component, attention, healthy, scale, patterns, positive, regional, sn, local, fluctuations, rsns, strength, analyses, altered, salience, control, global, rsfc |

Supplementary Table 2: Correct topics for all tasks in the 400 topic-set

| Task | Topic | Top Terms | |
|---|---|---|---|
| **Math** | 107 | 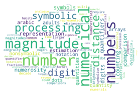 | number, numerical, numbers, magnitude, processing, distance, symbolic, digit, representation, counting, numerosity, comparison, digits, quantifiers, symbols, nonsymbolic, estimation, arabic, quantity, dots, arrays, quantities, adults, notation, magnitudes, dyscalculia, abstract, numerals, decade, numerosities, sulci, larger, represented, congruity, dot, comparisons, quantifier, line, format, fractions |
| | 113 | 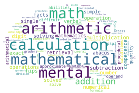 | arithmetic, calculation, mathematical, mental, math, addition, multiplication, subtraction, retrieval, operations, operation, competence, number, numerical, solving, single, verbal, hips, digit, solved, facts, exact, mathematics, simple, abacus, approximate, arithmetical, abilities, procedural, rely, operand, quantity, pspl, solve, difficult, competent, numeral, access, graders, formula |
| **Memo** | 014 | 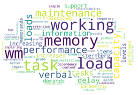 | wm, load, memory, working, task, maintenance, performance, capacity, loads, verbal, information, tasks, delay, increasing, increased, levels, items, support, sternberg, demands, parametric, performed, maintained, memoranda, automatization, resource, delayed, probe, match, contents, sample, probes, dependency, retention, letter, temporary, impair, epochs, demand, content |
| | 376 | 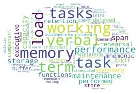 | memory, working, task, verbal, tasks, term, load, performance, maintenance, storage, capacity, span, performed, functions, executive, rehearsal, retention, mnemonic, sternberg, delayed, store, component, buffer, digit, maintaining, individual, stored, intra, capacities, demand, remember, seconds, subcomponents, hold, attenuated, memorizing, probe, inefficiency, rst, match |
| **Video** | 198 | 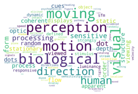 | motion, biological, moving, visual, perception, direction, human, processing, sensitive, form, dot, stationary, coherent, static, dots, process, flow, order, structure, optic, cues, viewing, random, depth, displays, apparent, viewed, responsive, sfm, object, luminance, dynamic, strongly, stimulus, defined, sensitivity, velocity, stream, selective, speed |
| **Rest** | 013 | 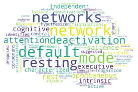 | network, default, mode, networks, resting, deactivation, attention, spontaneous, cognitive, rest, salience, executive, independent, intrinsic, characterized, fluctuations, scale, active, component, central, dmn, cognition, hypothesized, identified, suggested, proposed, deactivations, failure, shown, absence, consistent, recent, elements, anticorrelated, intrinsically, anti, functionally, deactivate, examination, referential |
| | 233 | 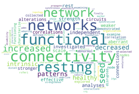 | connectivity, functional, networks, resting, network, increased, seed, patterns, decreased, intrinsic, healthy, independent, analyses, rest, investigated, scale, strength, correlations, alterations, stronger, effective, connections, weaker, circuits, aberrant, demonstrated, nodes, examine, correlation, characterize, collected, functioning, explored, node, disrupted, abnormal, organization, selected, rsfmri, examining |
| | 337 | 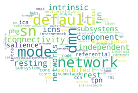 | dmn, network, default, mode, sn, connectivity, resting, independent, component, networks, rest, icns, intrinsic, subsystems, fpn, tpn, salience, exhibited, phg, referential, ica, subsystem, core, ccn, smn, intra, mental, anticorrelation, node, subnetworks, rai, internal, tcn, anticorrelations, inter, vmax, coherent, internetwork, hubs, edp |

Decoding accuracy was evaluated quantitatively using the ranking accuracy (*RA*) metric (15), which is defined as follows for a given subject (*s*) and task (*t*):

$$RA_{s,t} = 1 - \frac{RankCorrect_t - 1}{Ntopics - 1}$$

where *Ntopics* is the number of topics in the set (e.g., 50 or 400), and $RankCorrect_t$ is the decoding rank for the correct topic. The p-value for each median rank accuracy score across subjects was obtained using draws from the distribution under the null hypothesis of chance performance (uniform rank position). Given a sample of null results for each subject, we computed the median across subjects, to yield a null distribution for the median score. The p-value for the actual rank accuracy score was the probability mass in the tail for values greater than or equal to that score. When there is more than one correct topic, $RankCorrect_t$ refers to the average decoding rank across all those correct topics.

#### Visualization of Windowed Connectivity

Windowed connectivity matrices, following vectorization contain thousands of elements, and consequently are hard to visualize; even with high-dimensional visualization tools such as parallel coordinate plots (16) or star glyphs (17). In such high dimensional scenarios, an alternative for aiding with visualization is to use manifold learning to bring the data into a 2 or 3-dimensional spaces while preserving, as much as possible, the structure of the data in its native high-dimensional space. Here, we use one particular manifold learning technique, Laplacian Eigenmaps (*LE*; (18)), to help interpret results and compare between rest and task conditions. Laplacian embeddings were chosen because of their computational tractability, their interpretational value from a clustering perspective, and because they place emphasis on the preservation of local geometry in the higher dimensional originating space.

Laplacian eigenmaps were computed with the *scikit-learn Python* library (19). In all instances we used correlation distance as the dissimilarity metric, to be consistent with the *k-means* portion of the study. Input affinity matrixes to the Laplacian eigenmap algorithm were constructed using the nearest neighbor approach (NN=100).

### Supplementary Figures

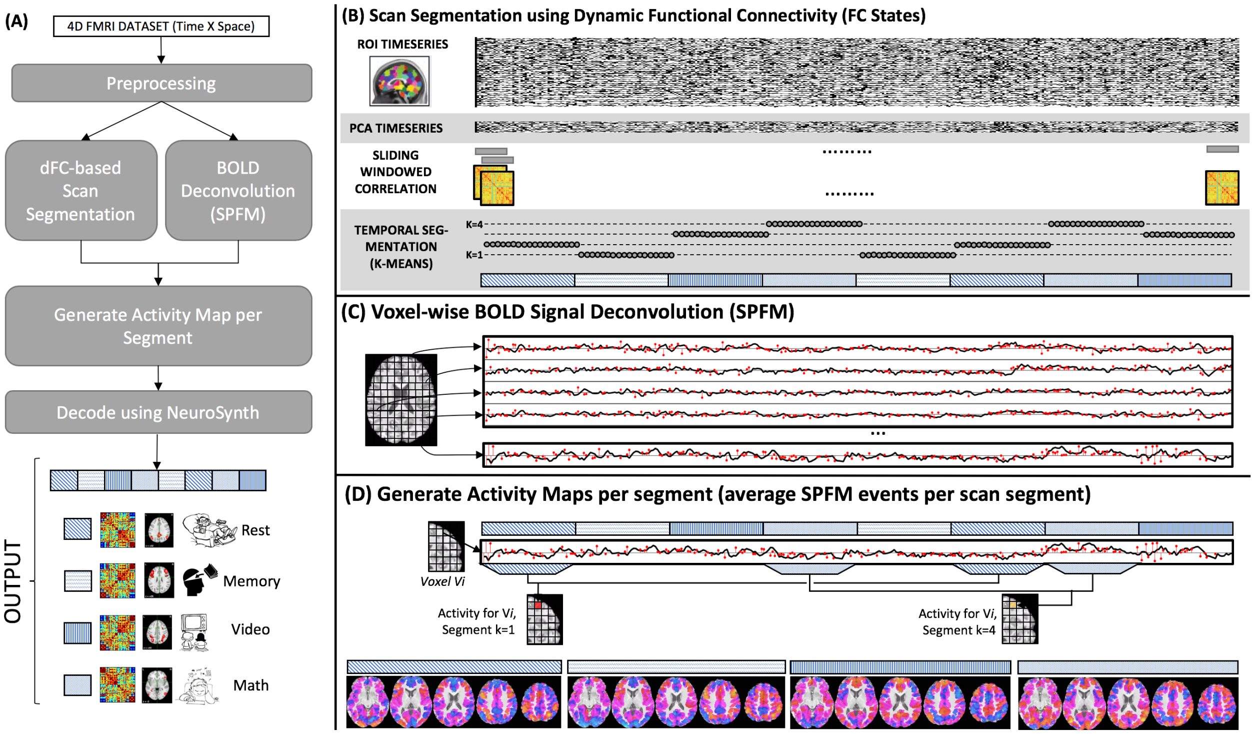


Supplementary Figure 1. Schematic of the analysis pipeline. (A) High level start-to-end schematic of analyses for the segmentation and decoding of task periods in the multi-task dataset. (B) Depiction of the main steps involved in the scan segmentation portion of the analyses. Following pre-processing, representative timeseries were obtained for all ROIs and those were input to a PCA step to reduce the dimensionality of the data. The next step was to compute sliding windowed correlation matrixes that were subsequently entered to a k-means analysis to assign each window to one of four segments. (C) Deconvolution was performed at the voxel-wise level. For each voxel, the deconvolution algorithm takes as input a pre-processed timeseries (black curve) and produces as output a time series of sparse events (red stems). (D) Activation maps per segment were generating by averaging event time series (red) within the confines of each segment as determined by the k-means step (blocks with same filling patterns). After completing this operation of each voxel, a full brain activity map was obtained per scan segment (bottom of panel D)

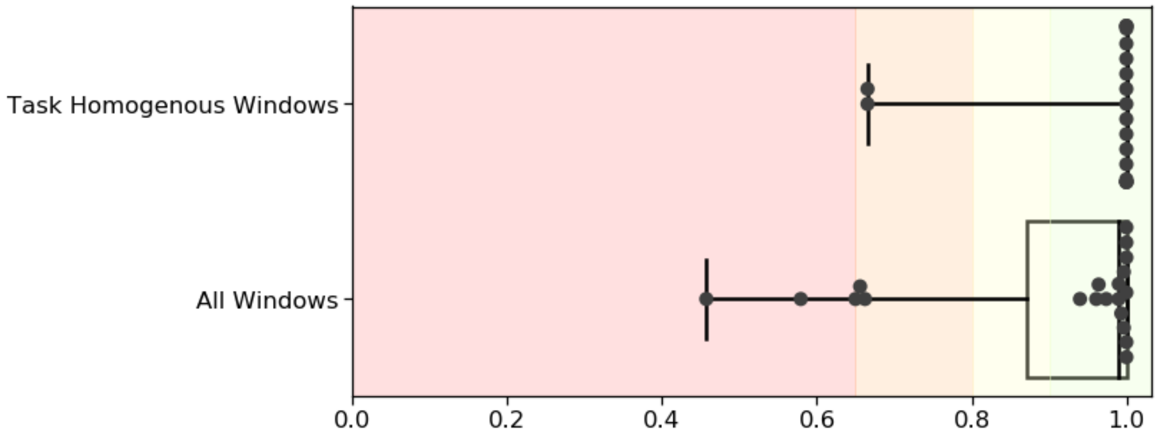


Supplementary Figure 2. Group-level quantitative evaluation of how well FC-states captures the task structure of the multi-task scans. Top boxplot shows results when only task-homogenous windows (e.g., those that fall completely within the temporal span of individual task blocks) enter the analysis. The bottom boxplot shows results when all available overlapping windows enter the analysis. Background colors depict common interpretational levels for ARI: <0.65 poor recovery of original structure; 0.65 < ARI < 0.8 moderate recovery; 0.8 < ARI < 0.9 good recovery; and ARI > 0.9 perfect recovery. In each boxplot, black dots represent different subjects, the boxed area shows the quartiles of the data, and the whiskers extent to show all remaining non-outlier data points.

#### 50 TOPICS SET – Topics clearly associated with the four tasks of interest

| Task | NeuroSynth Topic ID | NeuroSynth Reverse Inference Map |
|---|---|---|
| **Math** | 042 | 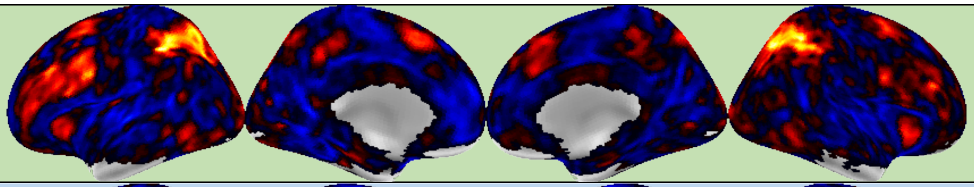 |
| **Memo** | 022 | 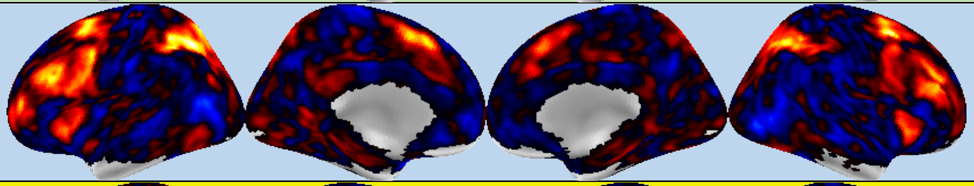 |
| **Vide** | 011 | 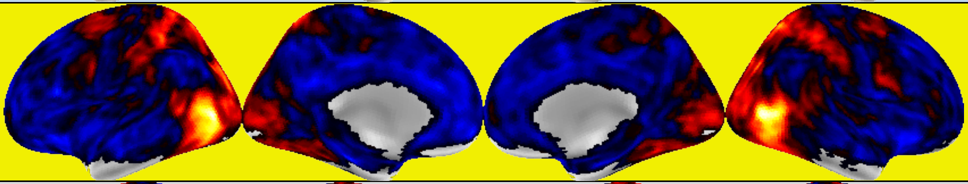 |
| **Rest** | 010 | 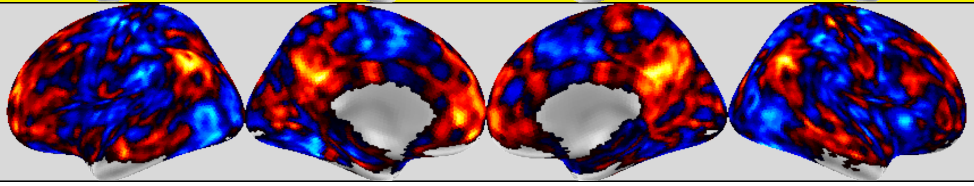 |

Supplementary Figure 3. Meta-analytical reverse inference maps for the "correct" topics from the 50-topic set associated with each of the tasks performed during the multi-task scans. The Topic ID corresponds to the numerical identifier of the topic in the NeuroSynth platform.

#### 400 TOPICS SET – Topics clearly associated with the four tasks of interest

| Task | NeuroSynth Topic ID | NeuroSynth Reverse Inference Map |
|---|---|---|
| **Math** | 107 | 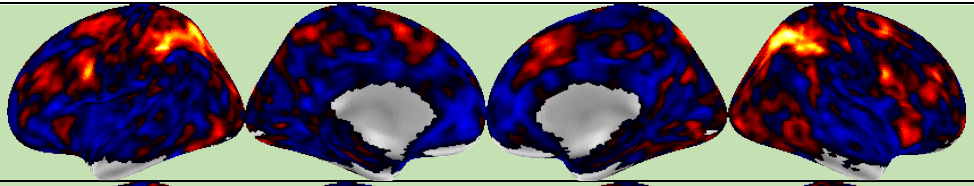 |
| | 113 | 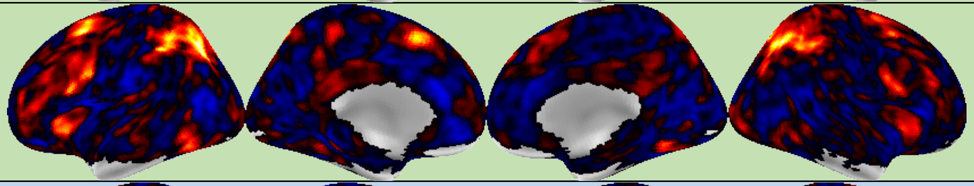 |
| **Memo** | 014 | 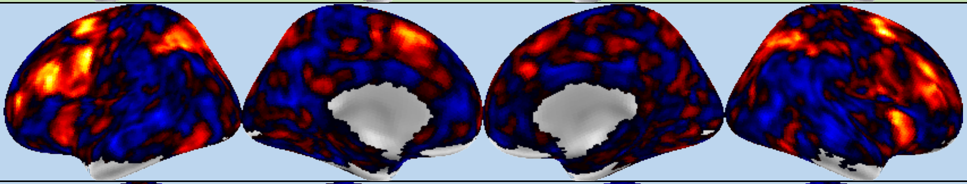 |
| | 376 | 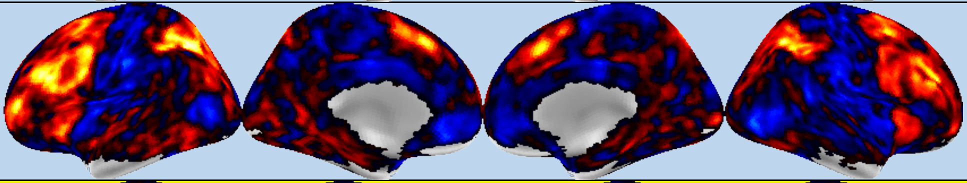 |
| **Vide** | 011 | 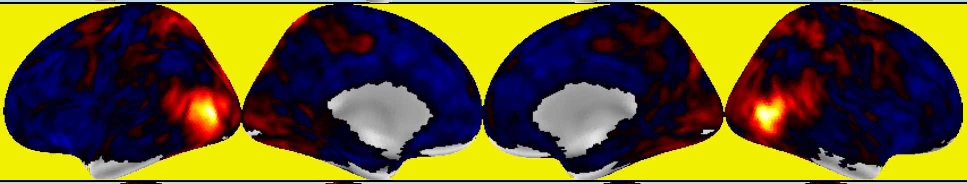 |
| **Rest** | 013 | 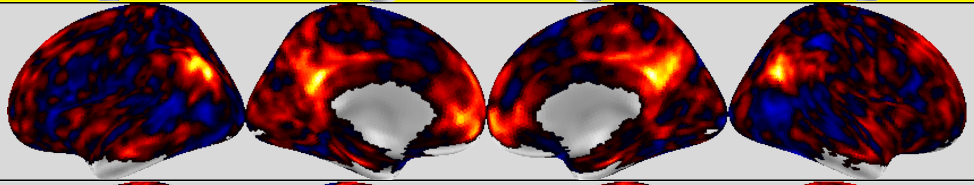 |
| | 233 | 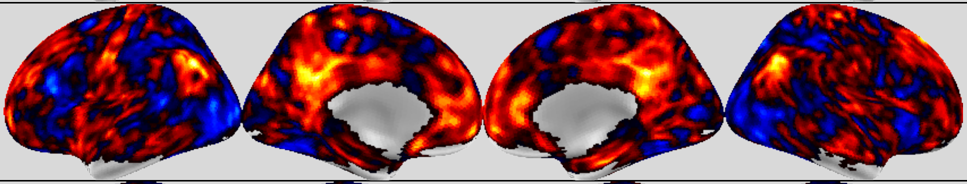 |
| | 337 | 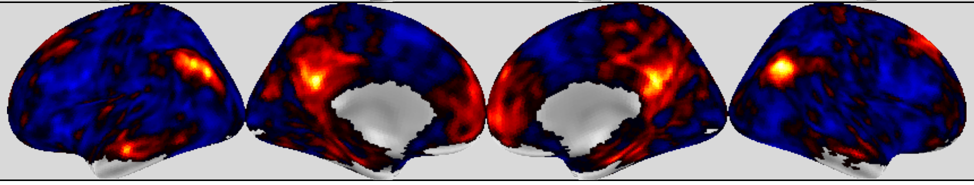 |

<u>Supplementary Figure 4</u>. Meta-analytical reverse inference maps for the “correct” topics from the 400-topic set associated with each of the tasks performed during the multi-task scans. The Topic ID corresponds to the numerical identifier of the topic in the NeuroSynth platform.

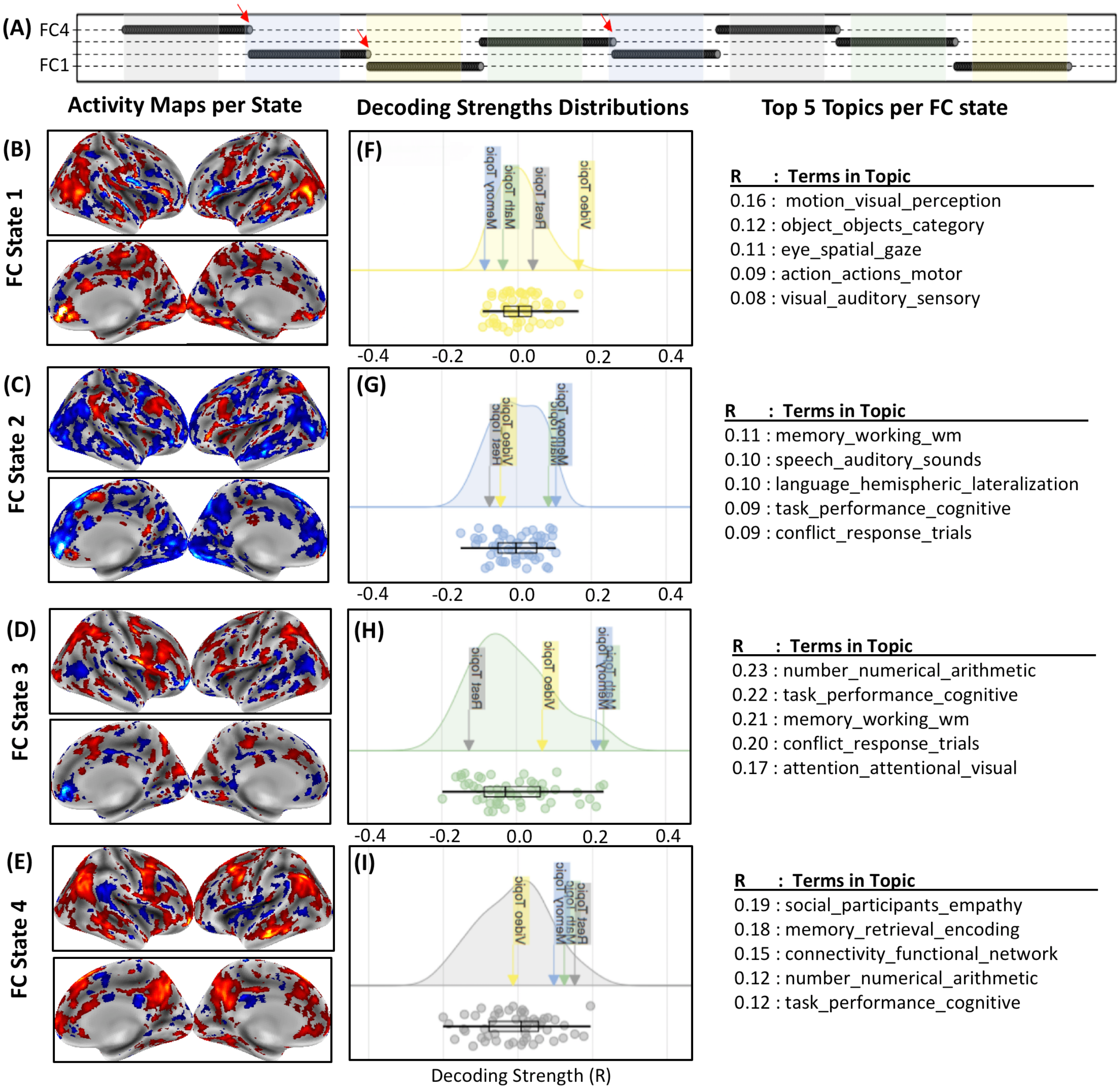


Supplementary Figure 5. Individual subject task decoding results for the 50-topics set. (A) Scan segmentation results. Each dot represents a snapshot of windowed connectivity. The position of the dot in the y-axis indicates to which FC-state the snapshot of windowed connectivity was assigned. Tasks periods are depicted as colored bands for reference (REST: gray; MEMO: blue; MATH: green; VIDEO: yellow). (B-E) Activity maps for each FC-state obtained with SPFM. Red arrows indicate small segmentation mistakes for windows in the vicinity of task blocks onsets. (F-I) Decoding results in the form of cloud plots and top-five lists. Cloud plots depict the probability distribution of decoding strength values across all 50 topics for each FC state in the form of kernel density estimates (colored curves), swarm plots (colored dots; one per topic) and boxplots (black). In these plots the location of "correct" topics per task are clearly marked by boxed text with arrows. Finally, the tables on the right of the figure lists the top 5 topics with the highest decoding strength for each FC state. Topic names are constructed using the top three terms associated with the topic.

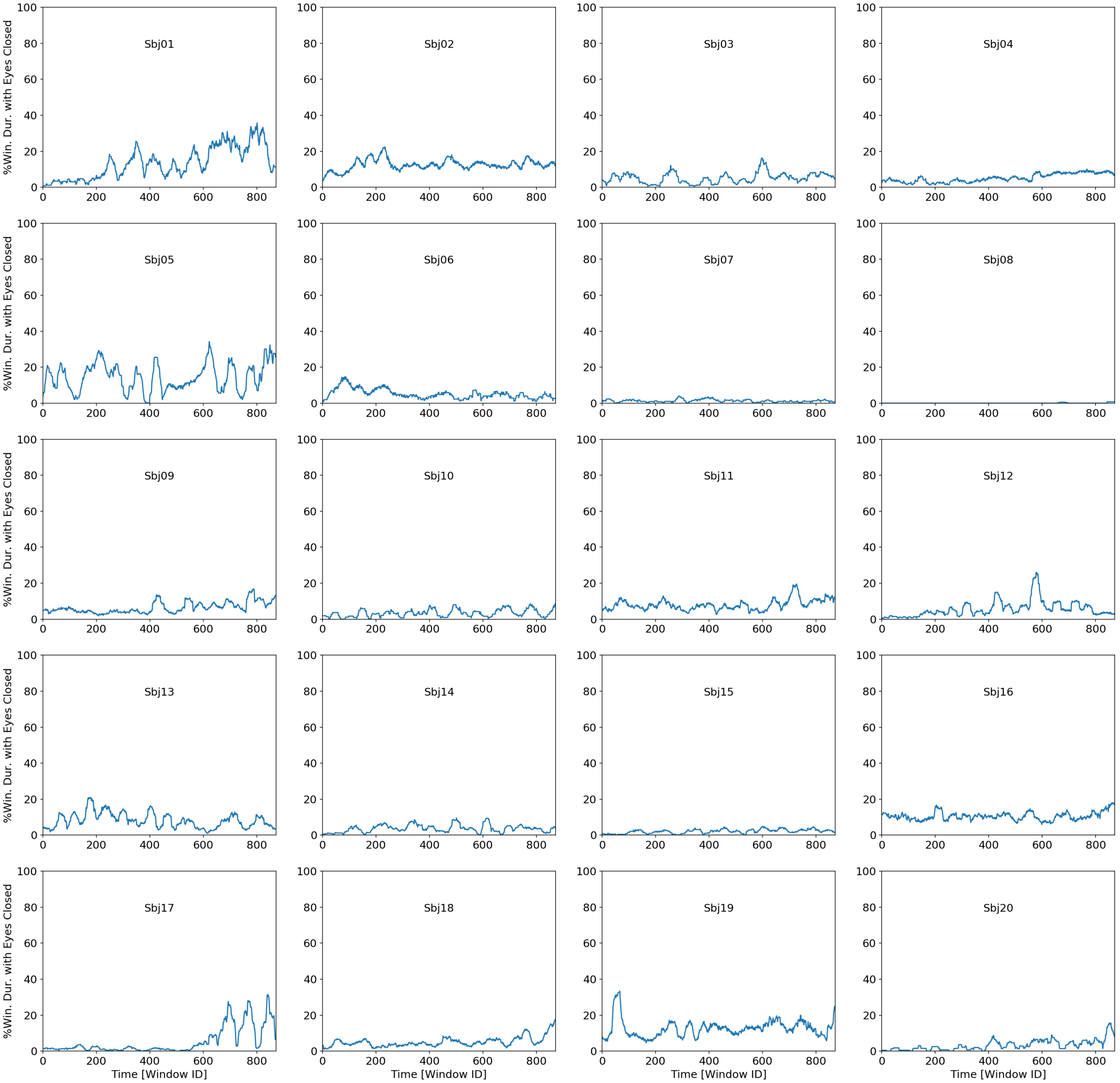


Supplementary Figure 6. Traces of percentage of time within each window for which subjects had their eyes closed according to the concurrent eye tracking recordings made available by the HCP consortium. For all subjects, time of eye closure was never above 40% in any single window, signaling that subjects remained awake for the duration of the scans.

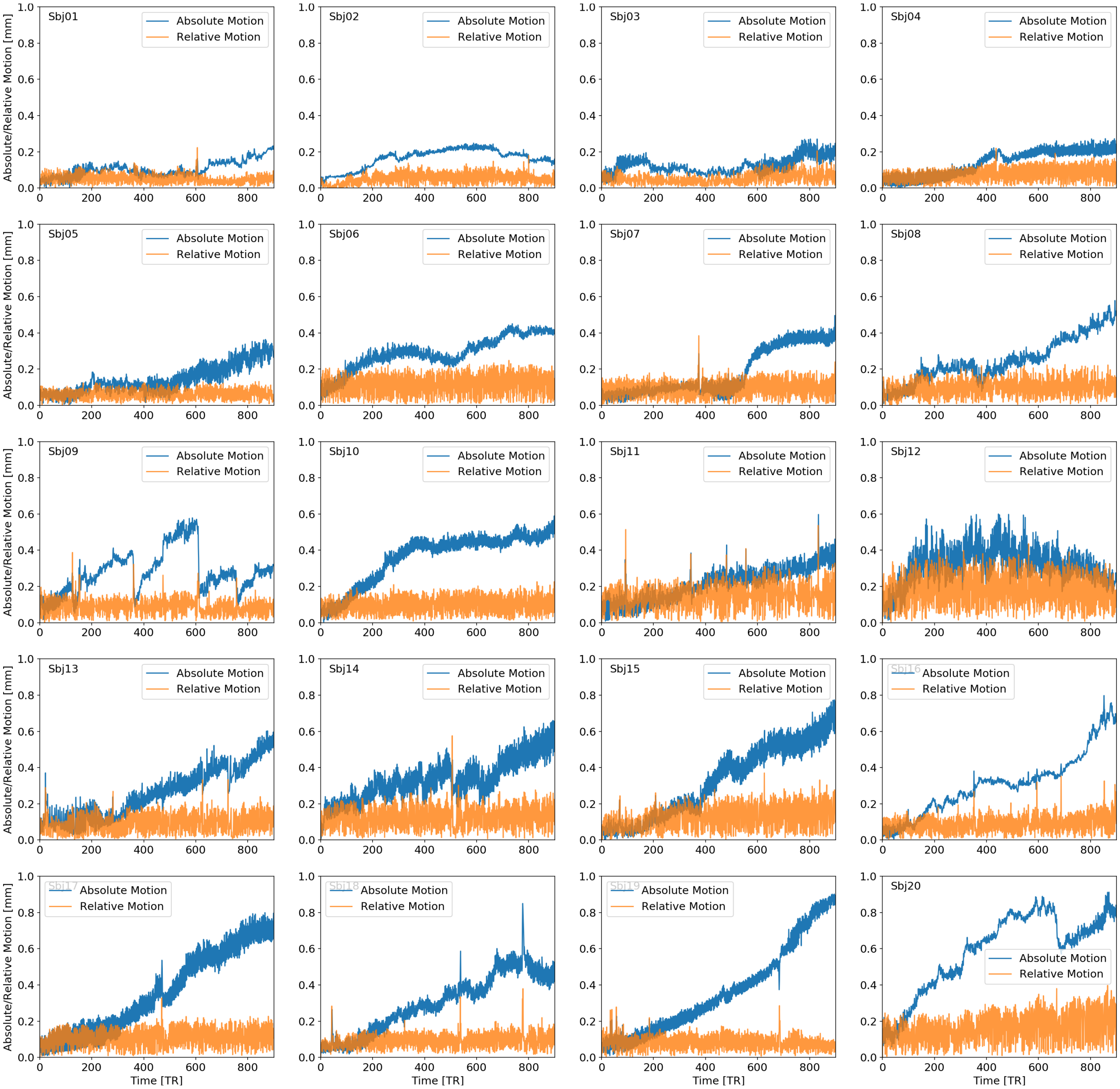


Supplementary Figure 7. Traces both absolute (blue) and relative volume-to-volume (orange) motion in mm for all selected subjects from the HCP dataset.

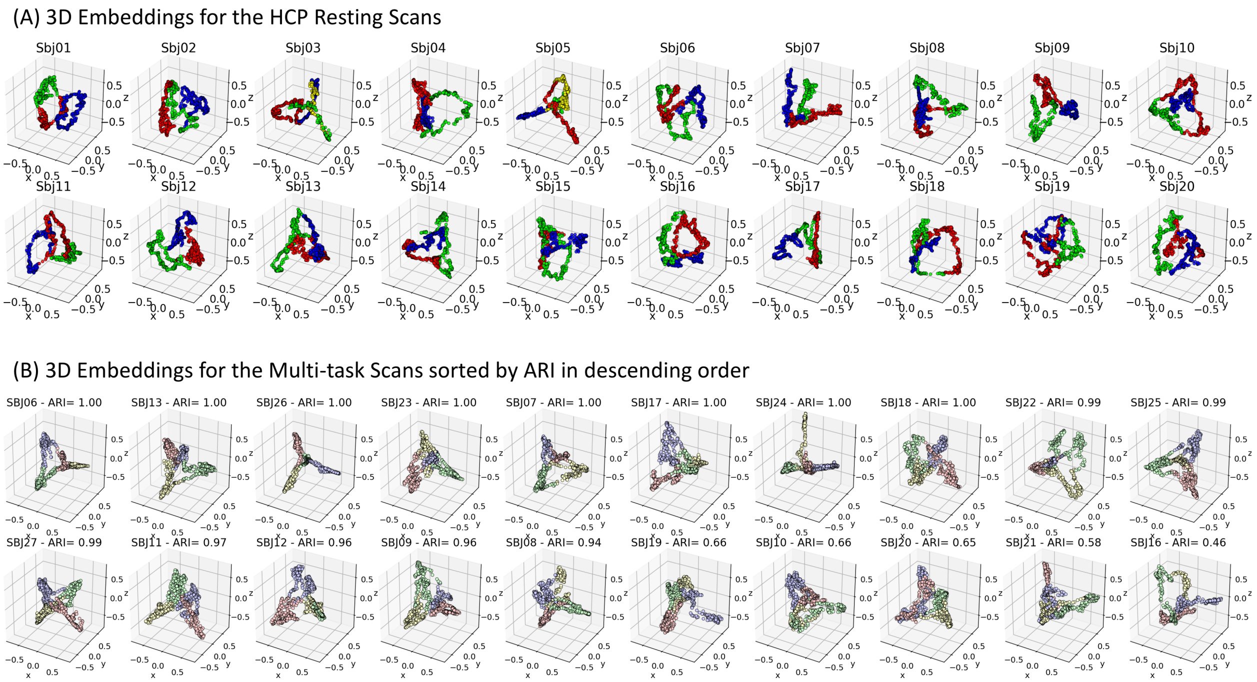


<u>Supplementary Figure 8</u>. (A) 3D embeddings for all 20 resting subjects from the HCP dataset. Connectivity snapshots are colored according to automatically detected FC states. Only for two subjects (sbj03 and sbj05) the algorithm detected 4 FC-states. For all other subjects, the number of states equal 3. (B) 3D embeddings for all 20 subjects in the multi-task dataset. Snapshots are colored according to FC state. Subjects in panel B are sorted according to ARI in descending order.

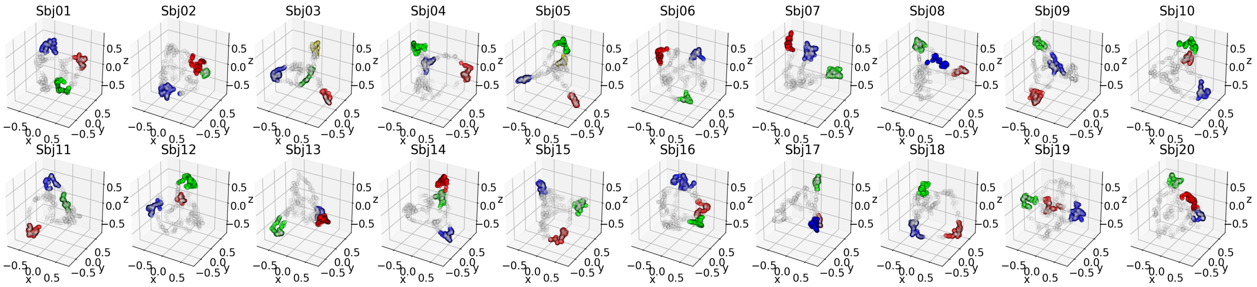


<u>Supplementary Figure 9</u>. Same embeddings as in supplementary figure 8.A (pure rest scans), but this time only windows detected to sit near the distal ends of the embeddings are colored.

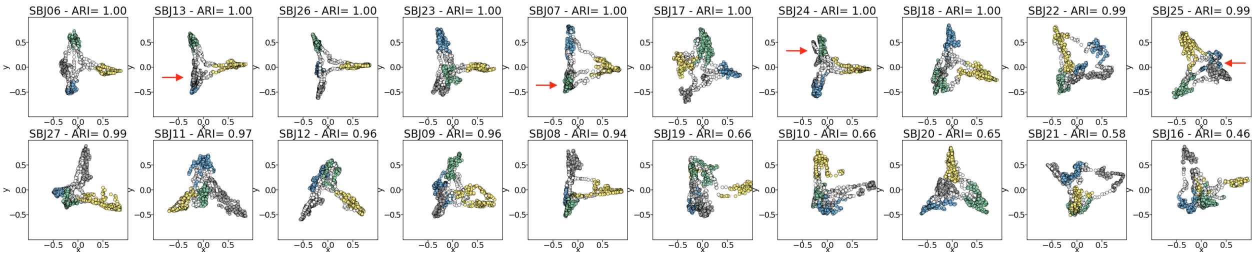


<u>Supplementary Figure 10</u>. Two dimensional Laplacian Embeddings for all subjects in the multi-task dataset. Connectivity snapshots are colored by task (grey = rest, blue = memo, green = math, yellow = video, white = task inhomogeneous windows). Red arrow signal occurrences where the excessively low dimensionality of the 2D space lead to crowding of windows from distinct tasks into a single location of space in subjects with high ARI.
